# CurateMake: an auditable workflow for multi-source ITS reference database harmonisation and phylogenetic validation

**DOI:** 10.64898/2026.07.10.737676

**Authors:** Auguste Gardette, Eugeni Belda, Edi Prifti, Jean-Daniel Zucker

## Abstract

1. Reference databases shape the taxonomic resolution, uncertainty, and reproducibility of metabarcoding analyses. For ITS barcodes, public references are distributed across repositories with different taxonomic conventions, geographic coverage, and annotation practices, creating conflicts, missing ranks, and misannotations when databases are merged or compared.
2. We introduce **CurateMake**, a reproducible Snakemake workflow for ITS reference database construction, harmonisation, and validation. It integrates four public sources (UNITE, BOLD, PLANiTS, and CALeDNA) and user-supplied databases, combines Catalogue of Life name harmonisation with ITSx-based region standardisation, MSA/HMM-based alignment grouping, and SATIVA phylogenetic validation. Raw, CoL-harmonised, and SATIVA-validated annotation layers are retained throughout to compare curation effects while preserving flagged records for review.
3. We evaluated **CurateMake** on 3.58 million ingested sequences and controlled error-injection simulations. ITSx expanded the final harmonised database to 5.19 million barcode-resolved entries by recovering ITS1 and ITS2 sub-regions from full-length ITS records. Across the full dataset, normalised intra-cluster entropy decreased from Raw to CoL-harmonised to SATIVA-validated annotations, consistent with improved taxonomic coherence. In simulations, **CurateMake** achieved the highest correction rate across 1%–50% corruption and, at 15% corruption, corrected 42% ± 1% of introduced errors, compared with 28% ± 1% for CoL alone and 0% for SATIVA without the workflow’s alignment infrastructure.
4. These results show that nomenclatural harmonisation and phylogeny-informed validation address complementary error classes, with phylogenetic validation contributing measurably only within taxon-coherent alignments in this benchmark. **CurateMake** therefore provides a reproducible, provenance-tracked framework for auditable ITS reference database curation in metabarcoding workflows.

## 1 Introduction

Metabarcoding-based biodiversity assessments depend on the reference databases used to assign environmental sequences to taxa. These databases determine which taxa can be detected, the taxonomic resolution at which ecological patterns can be interpreted, the uncertainty propagated into community-level estimates, and the comparability of results across studies and time series. For ITS barcodes, this dependency is especially challenging because no single repository provides complete and uniformly curated coverage across the organisms for which ITS is used.

Public ITS reference sequences are assembled from independent curation initiatives that reflect distinct biological priorities, geographic emphases, and nomenclatural practices. UNITE was originally developed for fungal molecular identification and has expanded to include other eukaryotic sequences; PLANiTS targets vascular plant ITS; BOLD emphasises barcode-level data across curated taxonomic groups; and CALeDNA reflects the geographic and ecological priorities of California-focused biodiversity inventories (Abarenkov et al., 2024; Banchi et al., 2020; Ratnasingham et al., 2024; Meyer et al., 2021). Because these resources evolve independently, the same taxon may carry incompatible labels across sources, entire lineages may be absent from any individual database, and annotation errors, including misidentification, contamination, and nomenclatural drift, can propagate into taxonomic assignment and ecological interpretation (Cheng et al., 2023; Goudey et al., 2022; Keck et al., 2023), with direct consequences for applications such as detecting illegally traded species from plant ITS metabarcoding (Boer et al., 2017). Taxonomic concepts also shift through synonymies and nomenclatural updates that are not synchronised across repositories, introducing silent label changes that can compromise cross-study comparability (Patterson et al., 2016).

These inconsistencies have concrete consequences for downstream ecological analyses. In biomonitoring and invasion ecology, incomplete or nomenclaturally misaligned references can produce false detections or prevent early identification of target taxa when their sequences are absent or mislabelled (Jerde et al., 2011; Mugnai et al., 2023). In community ecology, uneven taxonomic resolution can affect estimates of beta diversity, temporal turnover, and community composition (Heino, 2014; Serrana et al., 2022; Lopes et al., 2021). Because ITS is used across fungi and plants, annotation errors may also propagate into cross-kingdom comparisons and functional guild assignments (Thomsen and Willerslev, 2015; Nguyen et al., 2016). A common workaround is to construct local or region-specific reference sets from existing repositories, but this approach cannot recover taxa absent from the selected source and may reinforce taxonomic or geographic gaps when databases evolve independently.

The bioinformatics landscape for ITS analysis provides well-developed solutions for read processing and classification, but fewer tools address reference database construction itself. QIIME2, DADA2, PIPITS, and ITSxpress support denoising, ITS region extraction, and taxonomic classification, but generally treat the reference database as a fixed input (Bolyen et al., 2019; Callahan et al., 2016; Gweon et al., 2015; Rivers et al., 2018). RESCRIPt extends QIIME2 with dereplication and taxonomy filtering within individual resources, but is not designed for integrated harmonisation across heterogeneous multi-source inputs (Robeson et al., 2021). SATIVA detects phylogenetically inconsistent labels within a given alignment, but requires taxon-coherent, pre-structured input and provides no mechanism for multi-source integration or nomenclatural standardisation (Kozlov et al., 2016). Recent semi-automated curation work on vascular plant ITS2 has demonstrated the value of combining sequence extraction with reference cleaning, but has not addressed cross-source harmonisation at scale (Quaresma et al., 2024). Existing tools therefore address complementary parts of the reference database problem, but, to our knowledge, no end-to-end framework currently integrates multi-source normalisation, nomenclatural standardisation, phylogenetic validation, and record-level provenance tracking within a single auditable workflow (Table 1).

**Table 1:** Comparison of **CurateMake** with related tools for ITS reference database preparation and curation. The comparison focuses on native support for features required in auditable multi-source reference database construction, not on the broader intended scope or quality of each tool.

| Tool | Multi-source schema<sup>a</sup> | Nomenclatural harmonisation | Phylogenetic validation | Reproducible workflow | Record-level provenance |
| --- | --- | --- | --- | --- | --- |
| **CurateMake** | Yes | Yes (CoL) | Yes (SATIVA) | Yes (**Snakemake**) | Yes |
| RESCRIPT | Limited | Partial | No | Yes (QIIME2) | Partial |
| PIPITS | No | No | No | Yes | Limited |
| ITSxpress | No | No | No | Yes | Limited |
| SATIVA | No | No | Yes | Limited | Limited |

<sup>a</sup> Multi-source schema refers to simultaneous ingestion, normalisation, and record-level tracking of heterogeneous reference sources within a shared data model. The terms 'No' and 'Limited' refer only to native support for the features listed in this table in the context of auditable multi-source reference database construction. They do not imply limitations in the tools' intended use cases or in their performance for read processing, classification, single-source database management, or phylogenetic label validation.

Here we present **CurateMake**, a reproducible Snakemake workflow for multi-source ITS reference database construction and curation. The workflow ingests and normalises sequences from UNITE, BOLD, PLANiTS, and CALeDNA, with support for user-supplied databases, and applies two complementary curation layers. The first layer harmonises nomenclature against the Catalogue of Life (Bánki et al., 2026), standardising taxonomic labels across sources while preserving unresolved records. The second layer applies SATIVA phylogenetic validation (Kozlov et al., 2016), identifying records whose labels are inconsistent with their phylogenetic placement. These layers are connected by an alignment-engineering step that includes ITSx extraction, hierarchical multiple sequence alignment construction with MAFFT, trimming, and HMM-based grouping into taxon-coherent alignment groups. Raw, CoL-harmonised, and SATIVA-validated annotation layers are preserved in parallel throughout the workflow, enabling direct quantification of the effect of each curation layer while retaining flagged and unresolved records for curator review.

We evaluate **CurateMake** through two complementary benchmarks targeting different dimensions of reference database quality. Entropy-based diagnostics quantify changes in label consistency within sequence clusters across the three annotation layers on the full dataset of 3.58 million ingested records. Controlled error-injection simulations with traceable ground truth quantify correction and recovery rates by error type and corruption level, providing a known-error benchmark for each curation layer independently and in combination. Together, these analyses test whether nomenclatural harmonisation and phylogeny-informed validation address complementary error classes, and whether the phylogenetic layer makes a measurable contribution only when taxonomic labels are evaluated within coherent multiple sequence alignments.

In this study, we aim to:

(1) provide a reproducible, auditable framework for harmonising multiple ITS reference databases into a consistent and provenance-tracked reference collection;
(2) quantify how nomenclatural harmonisation and phylogeny-informed validation affect taxonomic coherence using entropy-based diagnostics;
(3) evaluate correction performance under controlled conditions via error-injection simulations with traceable ground truth, reporting correction and recovery rates by error type and corruption level.

## 2 Methods

### 2.1 Data sources and versions

We assembled a full ITS reference dataset from four public sources: UNITE, BOLD, PLANiTS, and CALeDNA (Abarenkov et al., 2024; Ratnasingham et al., 2024; Banchi et al., 2020; Meyer et al., 2021). Datasets were extracted in late August 2025, with source release dates and retained record counts reported in Table 2. After ingestion and pipeline filtering, the full dataset contained 3,581,943 source records; a 4,842-record public mini-dataset was also generated for demonstration, testing, and runtime benchmarking. Counts are reported after pipeline-level filters. Barcode-level counts for ITS, ITS1, and ITS2 are provided in Table SS1; because parsed barcode records may count more than one barcode per source record, they should not be interpreted as source-record totals.

**Table 2:** Source databases used in this study. Counts correspond to records retained after ingestion and pipeline filtering.

| Database | Extraction date | Public release date | Full dataset | Mini-dataset |
| --- | --- | --- | --- | --- |
| UNITE | 21 Aug 2025 | 19 Feb 2025 | 2,069,189 | 1,034 |
| BOLD | 22 Aug 2025 | 22 Aug 2025 | 631,522 | 1,509 |
| PLANITS | 22 Aug 2025 | 29 Mar 2020 | 310,722 | 1,385 |
| CALeDNA | 29 Aug 2025 | 29 Aug 2025 | 570,510 | 914 |
| Total | — | — | 3,581,943 | 4,842 |

### 2.2 Workflow architecture

**CurateMake** comprises five sequential stages: multi-source ingestion, Catalogue of Life harmonisation, ITSx region extraction and deduplication, hierarchical MSA/HMM-based alignment assignment, and SATIVA phylogenetic validation (Figure 1). Each stage writes intermediate artefacts to a relational database and emits structured reports, preserving traceability from source records to final curated entries. Taxonomic annotations are modified only during CoL harmonisation and SATIVA-based validation; the intervening stages standardise barcode regions, remove redundancy, construct alignments, and assign sequences to alignment groups.

**Figure 1:**
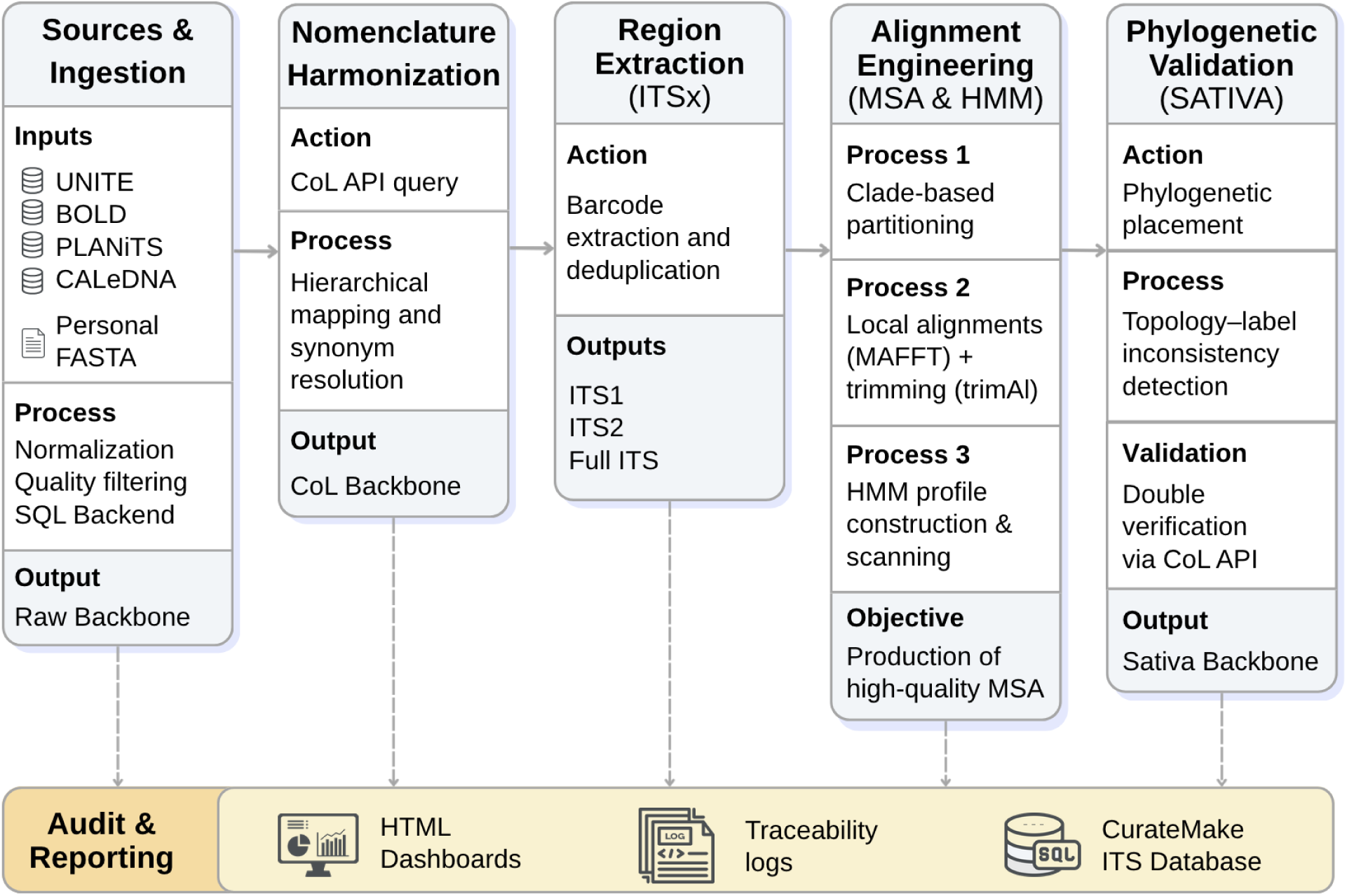
**CurateMake** pipeline overview. The workflow integrates source ingestion, CoL-based nomen-clatural harmonisation, ITSx region extraction, MSA/HMM-based alignment engineering, and SATIVA phylogenetic validation. These stages generate three parallel taxonomic annotation layers, Raw, CoL-harmonised, and SATIVA-validated. The audit and reporting layer exports HTML dashboards, structured reports, traceability logs, and the final curated ITS reference database.

A central design feature is the preservation of three parallel **taxonomic annotation layers**: **Raw**, corresponding to source taxonomy; **CoL-harmonised**, corresponding to nomenclature aligned against the Catalogue of Life; and **SATIVA-validated**, corresponding to phylogeny-informed correction suggestions after CoL validation. Here, annotation layer refers to taxonomic label sets stored in the database, not to phylogenetic tree structures. Retaining these layers enables direct comparison of nomenclatural harmonisation and phylogeny-informed validation while preserving provenance for curator review.

### 2.3 Data ingestion and database backend

Source-specific parsers normalise sequences, metadata, barcode fields, and taxonomic labels into a shared relational schema. User-supplied reference collections are supported through a FASTA template with associated metadata and taxonomy fields. Sequence quality filters, including minimum length and ambiguity thresholds, are defined in the central configuration file.

A configurable backend supports both small local runs and large-scale execution, using SQLite and PostgreSQL, respectively. The database acts as the workflow state store: each stage reads from and writes to explicit tables, preserving links among source records, deduplicated sequences, extracted barcode regions, alignment groups, CoL harmonisation outputs, SATIVA validation outputs, and final curated records.

### 2.4 Catalogue of Life harmonisation

To standardise nomenclature across heterogeneous sources, taxonomies are harmonised against the Catalogue of Life (CoL) API (Bánki et al., 2026). CoL was prioritised because its Base Release derives from expert-curated global species checklists and its annual DOI-versioned releases support reproducible analyses. Unlike the GBIF Backbone, which is a synthetic management classification for GBIF-mediated resources, or NCBI Taxonomy, which is organised around organisms represented in sequence databases, CoL provides a stable checklist-oriented reference for cross-source name harmonisation.

Taxonomies in the Raw annotation layer are matched hierarchically from the most specific available rank to the broadest rank: species, genus, family, order, class, phylum, and kingdom. Each query combines the name at the current rank with available higher-rank context, and fuzzy matching is enabled to accommodate minor spelling, typographic, and formatting differences.

On successful match, the full CoL classification populates the CoL-harmonised annotation layer. When matching occurs at a higher rank, unresolved lower ranks are retained from the Raw layer to preserve source information. Sequences with no CoL match at any rank are flagged as COL_NO_MATCH and retain their Raw taxonomy for inspection.

### 2.5 ITSx extraction and post-processing

Sequences are deduplicated before ITSx to reduce redundant computation, using 100% identity over full-length sequences with identical associated taxonomy. Records are then split by barcode type and processed with ITSx (Bengtsson-Palme et al., 2013) to identify ITS, ITS1, and ITS2 regions. Outputs are parsed per barcode type to update sequence status and reinsert extracted regions into the database.

A second deduplication pass removes duplicates introduced by region extraction. These steps generate standardised barcode-level datasets, enlarge the effective reference collection by recovering ITS1 and ITS2 sub-regions from full-length ITS records, and preserve provenance links to the original source records.

### 2.6 Hierarchical MSA, trimming, and HMM-based assignment

#### 2.6.1 Task construction

MSA tasks are defined per barcode type and taxonomic group using a hierarchical manifest. Task construction starts at a configured taxonomic rank, family by default. Groups with fewer than 20 sequences are excluded because they provide limited phylogenetic context for label validation. Groups exceeding 1,000 sequences are recursively split to finer ranks, down to species; when no finer resolved rank is available, large groups are split into balanced sub-tasks. Tasks exceeding 2,000 sequences are flagged for manual inspection.

This strategy avoids validation units that are either too small to be informative or too large and heterogeneous to remain interpretable and computationally tractable. The task manifest provides a reproducible bridge between the taxonomic hierarchy and the alignment units used by SATIVA.

#### 2.6.2 Alignment and trimming

Each task is aligned with MAFFT using –auto and multithreading, allowing task-specific strategy selection (Katoh and Standley, 2013). Alignments are trimmed with trimAl using 75% minimum sequence overlap and 50% minimum residue occupancy (Capella-Gutiérrez et al., 2009). Trimming is conservative, aiming to remove poorly supported regions and identify incoherently aligned sequences rather than aggressively shorten alignments.

Sequences removed during trimming are not deleted from the database. They retain their Raw and CoL-harmonised annotations and can be re-attached to an alignment group through HMM-based assignment.

#### 2.6.3 HMM construction and classification

For each trimmed alignment, a DNA profile HMM is built with HMMER using hmmbuild –dna (Eddy, 2011). Profiles are merged first into barcode-level databases and then into a global HMM library. Before classification, low-information profiles are removed using two configurable hmmstat-based quality-control thresholds: mean relative entropy per position and the proportion of informative positions. In this study, profiles were retained only when both values were at least 0.2. These thresholds were used to reduce overly permissive hmmscan matches and are reported with other software parameters in Appendix C.

Sequences are matched in batches against the HMM library using hmmscan with an E-value threshold of 10^−3^. The top full-sequence hit passing this threshold assigns each sequence to a barcode type and alignment group; assignment reports retain the selected profile, E-value, and score to support auditing of multi-profile matches. This step updates barcode and alignment membership only, without modifying taxonomic annotations. Trimmed-out sequences are re-attached when they receive a valid match; unmatched sequences remain stored with their Raw and CoL-harmonised taxonomy but are excluded from SATIVA validation.

### 2.7 SATIVA validation and integration

SATIVA is executed on quality-trimmed taxon-level multiple sequence alignments to generate phylogeny-aware suggestions for taxonomic correction (Kozlov et al., 2016). Candidate mislabeled sequences are flagged through leave-one-out validation against maximum-likelihood reference trees inferred with RAxML (Stamatakis, 2014). To bound runtime, per-task limits are enforced: 20,000 sequences, 5,000 alignment positions, and 6 hours wall time. SATIVA is run in ultrafast mode with a confidence threshold of 0.4.

SATIVA suggestions are treated as hypotheses rather than definitive taxonomic truth. Each suggestion is validated against CoL using the same API parameters as in section 2.4 and stored with an explicit status code: `Correct` for CoL-validated suggestions, Check for suggestions requiring curator review, and `Failed` for suggestions that cannot be validated. This produces an auditable SATIVA-validated annotation layer without replacing the Raw or CoL-harmonised layers.

### 2.8 Reporting, reproducibility, and computational requirements

The pipeline emits structured JSON reports and HTML dashboards for major stages, including sequence flow, CoL matching, ITSx extraction, MSA analytics, HMM profile and assignment statistics, and SATIVA validation. Logs, benchmark files, and traceability tables link final curated records to source records, extracted regions, alignment groups, and taxonomic annotation layers.

All steps are orchestrated with `Snakemake` using rule-specific Conda environments and a centralised configuration file, enabling reproducibility across local and high-performance computing deployments (Mölder et al., 2025). Software versions and exact command-line parameters are reported in Appendix C (Table S15) and summarised in Table S16.

Runtime and resource usage scale with dataset size and alignment-task complexity. The public mini-dataset required approximately 1.5 hours on 55 CPUs, whereas the full dataset required approximately two months on 190 CPUs. ITSx and SATIVA were the dominant runtime bottlenecks, and memory pressure was driven mainly by MSA construction, hmmscan, and SATIVA validation.

### 2.9 Evaluation design

We evaluated **CurateMake** with two complementary experiments, jointly designed to characterise how taxonomic-label uncertainty propagates through the sequential stages of the workflow. Experiment 1 quantified intra-cluster taxonomic entropy across the Raw, CoL-harmonised, and SATIVA-validated annotation layers on the full reference dataset, where no complete ground truth is available. Because these three layers correspond to successive processing stages, comparing entropy across them measures how each stage propagates and, ideally, reduces label uncertainty at scale; it provides a large-scale proxy for changes in taxonomic coherence after nomenclatural harmonisation and phylogeny-informed validation. Experiment 2 used controlled error injection with fully traceable ground truth. Designed as a component ablation, it compared CoL alone, SATIVA alone, CoL-to-SATIVA, and the full pipeline to isolate the contributions of nomenclatural harmonisation, phylogenetic validation, and the alignment-engineering step linking them, and thereby to attribute error correction to specific stages of the sequential pipeline.

#### 2.9.1 Experiment 1: Intra-cluster taxonomic entropy

Sequences were clustered with MMseqs2 linclust at 95%, 97%, and 99% identity using –`cov-mode` 0, -c 0.8, and –`cluster-mode` 2 (Steinegger and Söding, 2017). Within each annotation layer, sequences without a taxonomy record were dropped, and missing or empty labels at any rank were assigned an explicit “Unknown” category rather than removed, so that annotation incompleteness itself contributes to intra-cluster heterogeneity. Clusters reduced to a single sequence were then excluded; this singleton filter is applied once on cluster membership and is therefore rank-independent, yielding a fixed set of non-singleton clusters per identity threshold (368,257, 400,109, and 451,211 clusters at 95%, 97%, and 99%, respectively). Pairwise layer comparisons used clusters with a defined entropy in both compared layers (matched clusters).

For each eligible non-singleton cluster, Shannon entropy was computed from taxonomic label frequencies from kingdom to species:

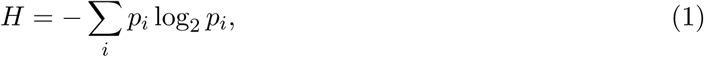

where *p_i_* is the relative frequency of the *i*-th label (including the “Unknown” category) within the cluster. Entropy equals zero for a taxonomically homogeneous cluster and increases with label diversity. To account for cluster-size variation, entropy was normalised by log_2_(*N*), where *N* is the number of sequences in the cluster (identical across ranks for a given cluster):

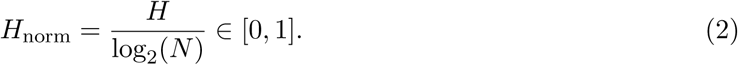

Mean normalised entropy was computed for each annotation layer, rank, and identity threshold. Pairwise differences between annotation layers were assessed with Wilcoxon signed-rank tests on matched cluster entropies, using significance thresholds of *p <* 0.05, *p <* 0.01, and *p <* 0.001. The primary visualisation reports mean normalised entropy ± standard error at the 99% identity threshold (figure 2); supplementary heatmaps show entropy across 95%, 97%, and 99% thresholds (Fig. SS1).

**Figure 2:**
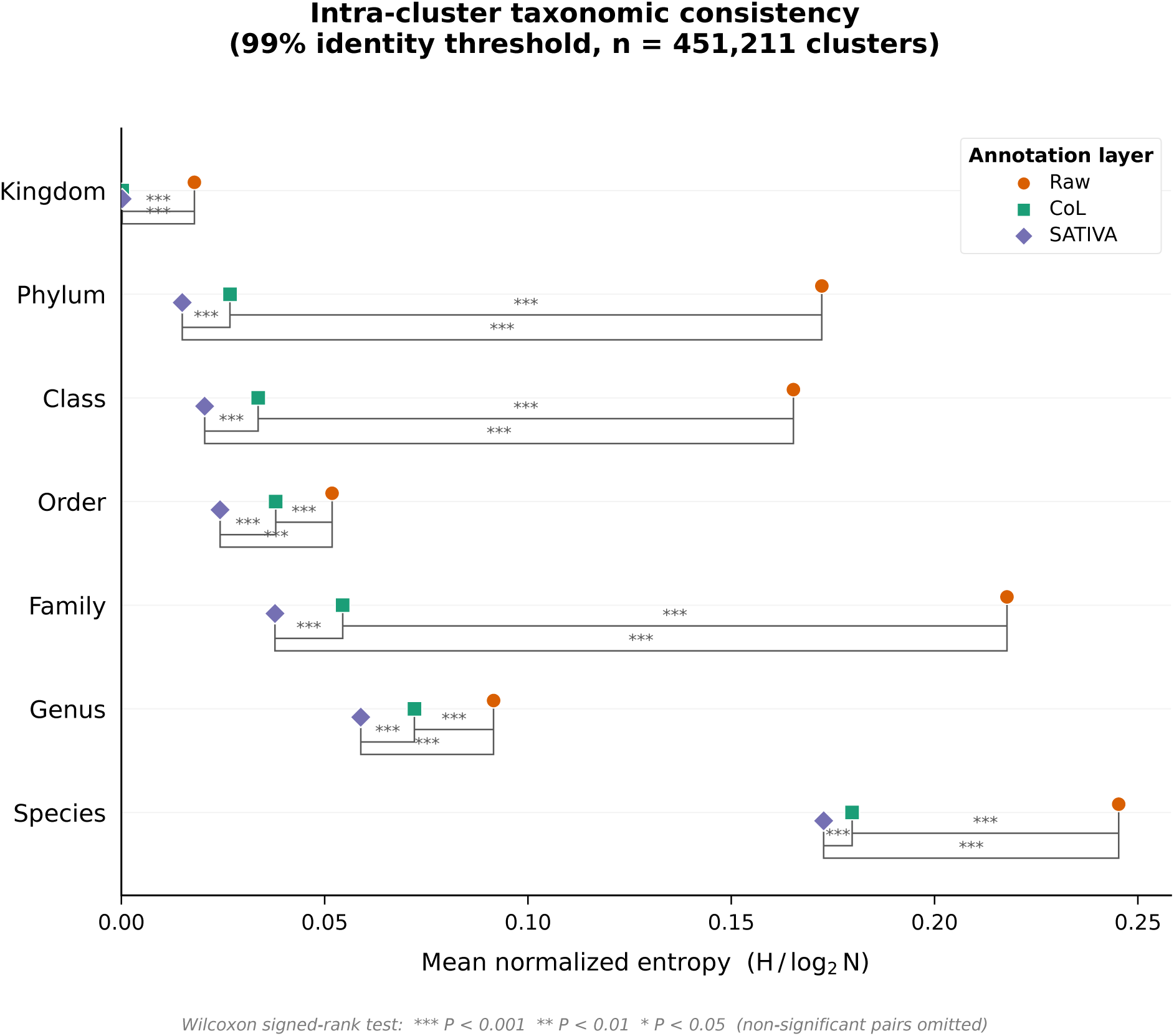
Mean normalised intra-cluster entropy (*H/* log_2_ *N*) per taxonomic rank and annotation layer at the 99% sequence identity threshold. Points represent means ± SEM across non-singleton clusters. Brackets indicate statistically significant pairwise differences between annotation layers (Wilcoxon signed-rank test; ^∗^*p <* 0.05, ^∗∗^*p <* 0.01, ^∗∗∗^*p <* 0.001; non-significant pairs omitted). The species rank exhibits the highest residual entropy across all three annotation layers. Analysis is restricted to non-singleton clusters (*n* = 451,211 at the 99% identity threshold).

#### 2.9.2 Experiment 2: Controlled error-injection simulation

##### Reference dataset construction

The simulation was designed as a controlled benchmark to compare correction strategies under fully traceable conditions, not to characterise taxon-wide ecological performance. Twenty full-ITS sequences were manually retrieved from BOLD in January 2026 and selected as ground-truth anchors: 10 plant species in *Carex* (Cyperaceae) and 10 fungal species in *Russula* (Russulaceae). These genera were chosen to provide complementary test contexts for cross-kingdom contamination and within-genus misidentification, while species selection was restricted to entries with CoL-accepted taxonomy and unambiguous BOLD records. This ensured that recovery could be evaluated against verifiable ground truth rather than confounded by nomenclatural ambiguity. The full anchor list, with BOLD process IDs and INSDC accessions, is given in Supplementary Table S2 (Appendix A.2).

The experimental dataset was built from Badread-simulated reads derived from these anchors. Synthetic sequences therefore do not represent the full biological diversity of either genus, but provide known taxonomies for strict output verification.

##### Sequence augmentation

Each of the 20 reference sequences was amplified in silico using Badread (Wick, 2019) with –identity 97,99,2. Read length was fixed to sequence length, with no junk reads or chimeras. After retaining only positive-strand reads to mimic amplicon directionality, the augmented dataset contained 1,989 sequences.

##### Taxonomy corruption

Corruptions were applied only to taxonomy fields; DNA sequences remained unchanged. For each scenario, a configurable proportion of sequences was sampled at random and assigned one of five error types with uniform probability by default: MISSING_RANK, OBSOLETE_TAXONOMY, MISIDENTIFICATION, CONTAMINATION, or UNIDENTIFIABLE. Corruption rates were 1%, 2%, 3%, 5%, 10%, 15%, 25%, and 50%, each run with five independent seeds, yielding 40 scenarios. All corruptions were traced in per-sequence reports recording original and corrupted taxonomies. Exact definitions of the five error types, the Badread augmentation parameters, and the four correction strategies are detailed in Appendix A.2.

##### Correction strategies

Because taxonomic annotations are updated only by CoL harmonisation and SATIVA integration, the simulation was structured as a component ablation. Each corrupted dataset was processed under four strategies: **Full pipeline**, combining CoL harmonisation, ITSx extraction, hierarchical MSA/HMM-based alignment assignment, and SATIVA validation; **SATIVA alone**, using kingdom-level MAFFT alignments and SATIVA without prior CoL harmonisation or the workflow’s hierarchical HMM-based alignment assignment; **CoL alone**, using Catalogue of Life harmonisation without phylogenetic validation; and **CoL-to-SATIVA**, applying CoL harmonisation followed by SATIVA without the full workflow’s hierarchical HMM-based alignment assignment. These conditions isolate the contributions of nomenclatural harmonisation, phylogenetic validation, and alignment engineering.

##### Metrics

Correction rates were first computed separately by error type. The per-error-type correction rate is the proportion of corrupted records of that type fully restored to ground-truth taxonomy, with all corrupted ranks corrected. For `CONTAMINATION`, full-rank correction was negligible, so partial detection was reported instead as the arithmetic mean of genus-level and kingdom-level recovery:

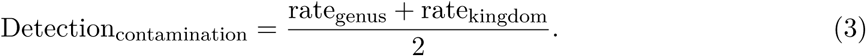

`UNIDENTIFIABLE` errors yielded null correction rates by construction because ranks replaced by Na provide no target label to recover; this error type is therefore excluded from both the per-error-type panel and the overall rate. The global (overall) correction rate was then defined as the unweighted mean of the four remaining per-error-type rates (`OBSOLETE_TAXONOMY, MISSING_RANK, CONTAMINATION` as partial detection, and `MISIDENTIFICATION`), computed per seed and then averaged across the five seeds; standard errors were likewise computed across seeds for each corruption rate.

## 3 Results

### 3.1 Quantitative results

The workflow expanded the reference collection from 3,581,943 ingested sequences to 5,189,645 unique, barcode-resolved entries in the final harmonised database. This increase was driven mainly by ITSx recovery of ITS1 and ITS2 sub-regions from full-length ITS records. After deduplication, the final database was approximately twice the size of the pre-ITSx dataset (2,536,799 sequences).

The taxonomic composition of the resulting database is summarised in table S6. Using the consolidated annotation per entry, defined as SATIVA where available, otherwise CoL, otherwise Raw, the database spans 7 kingdoms and approximately 183,000 species-level labels. Annotation completeness decreased towards finer ranks, with 46% of entries resolved to species overall and 75% within the plant subset.

#### 3.1.1 Taxonomic modification rates across sources

To quantify the scope of harmonisation, we measured, for each original sequence, whether curation introduced a nomenclatural conflict or a phylogenetic reassignment. Nomenclatural conflicts were defined as non-empty Raw ranks replaced by different non-empty CoL values, whereas phylogenetic reassignments were defined as non-empty CoL ranks replaced by different non-empty SATIVA values. These rates are conservative lower bounds because they capture detected differences between annotation layers rather than confirmed biological errors (table S11).

CoL harmonisation revealed both nomenclatural conflicts and rank completions. Conflict rates were moderate in UNITE and BOLD, affecting 9.9% and 13.4% of original sequences, respectively, but much higher in CALeDNA and PLANiTS because of source-specific kingdom-level labelling conventions. CALeDNA records are often labelled as “Eukaryota”, whereas CoL resolves them to kingdoms such as “Fungi” or “Plantae”; PLANiTS showed a similar convention-driven pattern.

Rank completions accounted for most Raw-to-CoL differences in UNITE and BOLD: CoL added information to previously empty ranks, without introducing conflicts, for 64.2% and 85.3% of sequences, respectively. SATIVA introduced fewer changes than CoL, with source-level phylogenetic reassignment rates ranging from 1.0% in UNITE to 9.5% in BOLD, but reached up to 12.5% among barcode-resolved ITS1 entries. At kingdom level, reassignment rates were lower in Fungi than in Plantae, reflecting both source composition and kingdom-labelling conventions.

#### 3.1.2 Experiment 1: Intra-cluster taxonomic consistency across annotation layers

At the 99% identity threshold (*n* = 451,211 non-singleton clusters), mean normalised intra-cluster entropy decreased monotonically from Raw to CoL to SATIVA at all taxonomic ranks (figure 2). CoL harmonisation alone significantly reduced entropy at every rank (Wilcoxon signed-rank test; *p <* 0.05 at kingdom, *p <* 0.001 from phylum to species), showing that nomenclatural standardisation resolves a substantial fraction of apparent label inconsistency within sequence clusters. SATIVA further reduced entropy after CoL harmonisation at most ranks, indicating an additional phylogeny-informed contribution after nomenclatural noise had been reduced. Complete mean entropies and pairwise Wilcoxon *p*-values for all ranks and identity thresholds are reported in Supplementary Table S5.

At kingdom level, entropy in the SATIVA-validated annotation layer reached near-zero (*<* 10^−3^), consistent with resolution of most high-level label conflicts. The CoL versus SATIVA pairwise difference at kingdom level was not significant and is omitted from figure 2, indicating that CoL alone resolves most kingdom-level inconsistencies.

Species-level entropy remained the highest across ranks even after the full workflow (Raw (≈ 0.245), CoL (≈ 0.180), SATIVA (≈ 0.173)), consistent with known limits of ITS for species-level discrimination (Schoch et al., 2012; Kiss, 2012) and, in part, with incomplete species-level annotation, which raises entropy independently of misannotation. Residual species-level entropy decreased with taxon abundance, and SATIVA’s additional reduction beyond CoL was concentrated among rare taxa (Supplementary Fig. S2). The same ordering of annotation layers was preserved across the 95%, 97%, and 99% identity thresholds (Supplementary Fig. S1), supporting the robustness of the comparison across clustering scales.

#### 3.1.3 Experiment 2: Controlled error simulation (component ablation)

Under fully traceable ground truth, the four correction strategies formed a component ablation: CoL alone tested nomenclatural harmonisation, SATIVA alone tested phylogenetic correction without prior CoL harmonisation, CoL-to-SATIVA tested sequential harmonisation and phylogenetic validation without the workflow’s MSA/HMM infrastructure, and the full pipeline tested SATIVA after workflow-level alignment engineering.

##### Overall correction rate

Across all corruption rates (1–50%), the full pipeline (**CurateMake**) consistently achieved the highest correction rate among the four strategies (figure 3, panel A). At 15% corruption, the reference level used for per-error-type comparisons, the full pipeline corrected 42.4% ± 1.2% of corrupted records (mean ± SEM, *n* = 5 seeds), compared with 27.5% ± 1.0% for CoL alone and 0.0% ± 0.0% for SATIVA alone. Across all strategies, the preservation rate for non-corrupted records was ≥ 99.8% at 15% corruption. Full numerical values for all corruption rates and strategies are given in Supplementary Tables S3 and S4.

**Figure 3:**
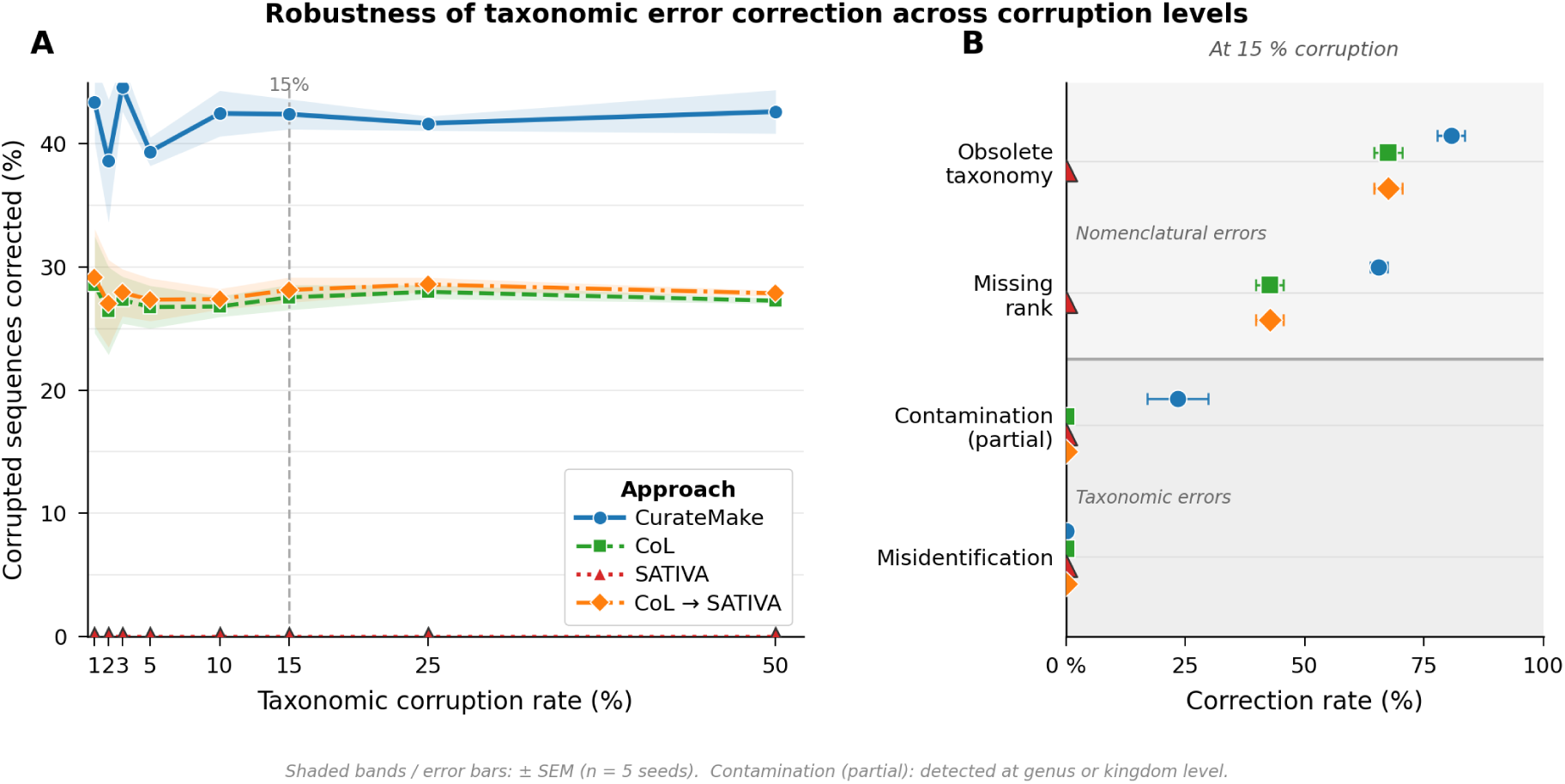
Controlled error simulation results. **(A)** Full-rank correction rate, defined as the proportion of corrupted sequences fully restored to ground-truth taxonomy, shown as mean ± SEM (*n* = 5 seeds) across taxonomic corruption rates from 1% to 50% for four correction strategies: **CurateMake** (full pipeline), CoL alone, SATIVA alone, and CoL-to-SATIVA without the pipeline’s MSA/HMM infrastructure. The dashed vertical line marks the 15% reference level. **(B)** Full-rank correction rate per error type at 15% corruption, except for Contamination (partial), which reports the mean of genus-level and kingdom-level recovery rates (see Methods). Unidentifiable errors, which yielded 0% correction across all strategies, are omitted.

SATIVA alone corrected no sequences across all tested corruption rates and seeds, and CoL-to-SATIVA performed similarly to CoL alone. Thus, sequential application of CoL and SATIVA was insufficient without the workflow’s hierarchical MSA/HMM-based alignment assignment. By contrast, the full pipeline exceeded CoL alone by approximately 15 percentage points at 15% corruption, showing that the alignment-engineering block is necessary for SATIVA to contribute in this benchmark.

##### Correction rate by error type

Correction capacity varied strongly across error types at 15% corruption (figure 3, panel B). The highest recovery rates were observed for `OBSOLETE_TAXONOMY` and `MISSING_RANK`: the full pipeline corrected 80.7% ± 2.9% and 65.5% ± 1.8% of these errors, respectively, compared with 67.5% ± 3.0% and 42.7% ± 2.9% for CoL alone. This indicates that CoL resolves many nomenclatural errors, while the full workflow adds recoveries through phylogeny-informed validation.

Full-rank correction of CONTAMINATION was negligible across strategies, but partial detection, defined as the mean of genus-level and kingdom-level recovery, reached 23.4% ± 6.4% for the full pipeline and 0.0% for CoL alone and SATIVA alone. No strategy meaningfully corrected MISIDENTIFICATION errors, and UNIDENTIFIABLE errors yielded null correction rates by construction because ranks replaced by Na provide no target label to recover.

##### Effect of corruption rate

The full pipeline remained the highest-performing strategy from 1% to 50% corruption (figure 3, panel A). Performance was broadly stable across moderate to high corruption levels, whereas uncertainty was larger at the lowest corruption rates because fewer corrupted records were available for evaluation.

#### 3.1.4 ITSx extraction, deduplication, and barcode assignment

Pipeline reports showed substantial reshaping of sequence flow during ITSx extraction, deduplication, and HMM-based barcode assignment (figure 4). ITSx was used to augment and standardise the reference collection by extracting ITS1 and ITS2 sub-regions from full-length ITS records, while retaining pre-existing barcode records for downstream assignment.

**Figure 4:**
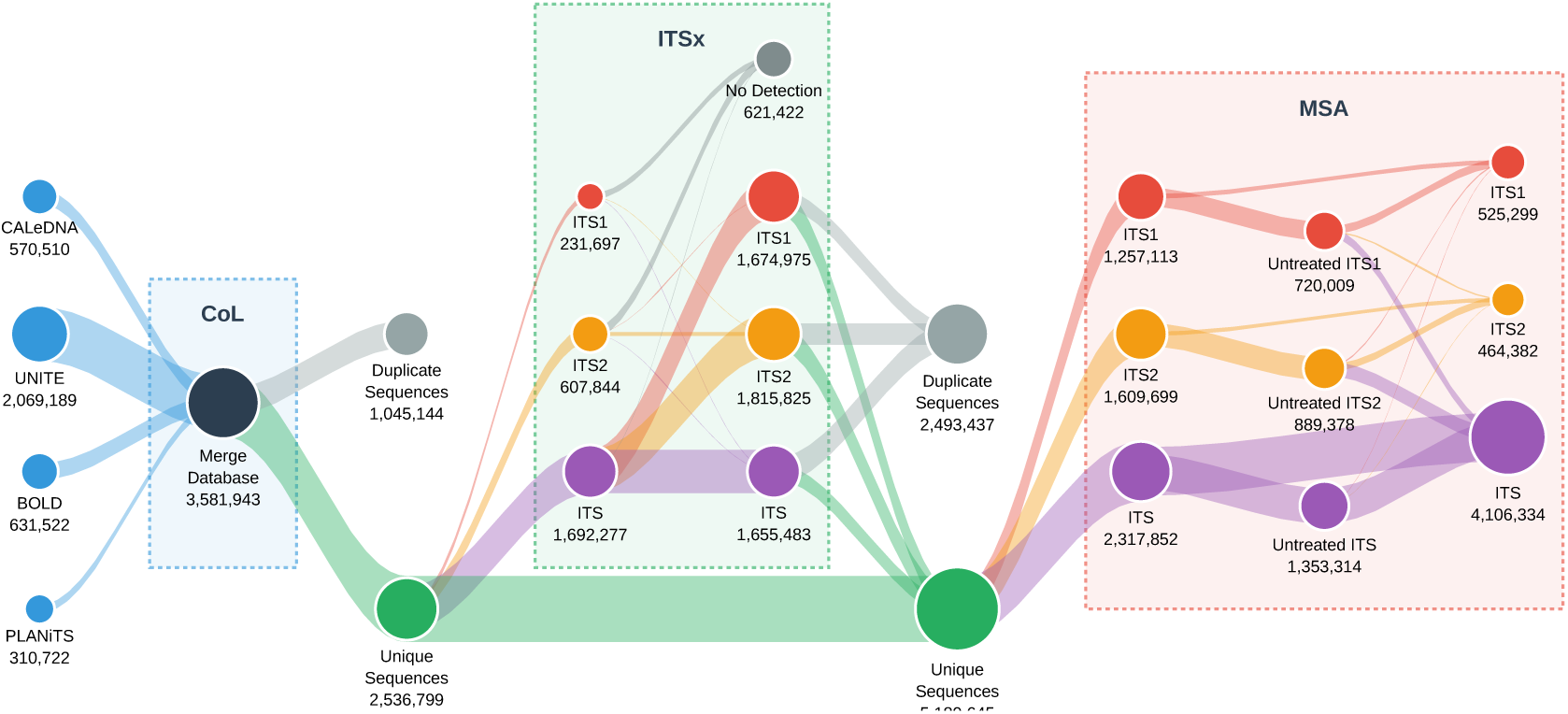
Sequence flow through the **CurateMake** pipeline on the full dataset after source-level filtering. Left panel: ingestion of retained records from the four source databases and CoL-based merging into a unified database, followed by an initial deduplication step. Centre panel: ITSx extraction, showing the number of ITS1, ITS2, and full ITS regions detected or extracted from each input barcode type, together with records for which no ITSx region was detected and duplicates removed after extraction. Right panel: sequence counts entering the MSA stage, split by assigned barcode group and ITSx treatment status, distinguishing extracted sub-regions from pass-through sequences. Colour coding indicates barcode group: ITS1, ITS2, and ITS.

After the first deduplication step, ∼2.54 million unique sequences were retained. Among these, ∼1.69 million full-length ITS records yielded ∼1.67 million ITS1 and ∼1.67 million ITS2 extracted sub-regions. Already trimmed ITS1 and ITS2 inputs produced few additional ITSx-derived entries because they often lack conserved flanking regions, but these original records were retained as pass-through sequences. After a second deduplication step removed ∼2.49 million duplicates introduced by sub-region extraction, the final harmonised database contained ∼5.19 million unique barcode-resolved entries. Numerical values for the extraction and deduplication steps are given in Supplementary Tables S12 and S13.

HMM-based assignment resolved an alignment group for ∼2.92 million sequences. Full-length ITS inputs almost always retained their ITS grouping label (99.9%), whereas many pre-trimmed sub-region inputs matched a full-ITS profile (48.8% of ITS1 and 63.9% of ITS2 inputs; table S14). This reflects inclusive profile matching during alignment assignment, not recovery of full-length sequence information. HMM assignment was therefore used only to organise sequences into barcode-specific alignment groups; downstream analyses retained the original ingestion barcode.

### 3.2 SATIVA suggestion patterns and qualitative examples

Representative SATIVA suggestions from the controlled simulation (seed 1, 15% corruption, full pipeline) are shown in table S7. They illustrate the main actionable classes observed in the benchmark, including kingdom-level contamination detection and missing-rank completions at order, family, and genus levels. Deletion suggestions are not emphasised because they remove annotation rather than replacing or completing it, and may discard biologically meaningful information in cases such as shared barcodes, hybridisation, or polyploidy.

On the full dataset, SATIVA produced ∼627,000 taxonomic suggestions across ∼18,250 alignment-defined taxonomic groups. Completions were the most frequent suggestion type (∼281,000; 44.8%), followed by deletions (∼174,000; 27.8%) and substitutions (∼172,000; 27.4%). We interpret completions and substitutions as the most directly actionable classes, whereas deletions are treated more cautiously because they remove existing annotation. The full breakdown by suggestion type, taxonomic rank, barcode, and source is given in Supplementary Tables S8, S9, and S10.

Each SATIVA suggestion is validated against CoL and stored in the SATIVA-validated annotation layer with an explicit status code (Correct, Check, or Failed). This allows curators to compare suggestions against the Raw and CoL-harmonised annotation layers rather than applying phylogenetic suggestions as definitive corrections.

## 4 Discussion

### 4.1 Summary of contributions

This work presents **CurateMake** as a reproducible and auditable framework for constructing taxonomic reference databases from heterogeneous sequence sources. Its central contribution is a layered curation architecture that separates source provenance, nomenclatural harmonisation, alignment engineering, phylogeny-guided validation, and database export into traceable steps. This makes reference database curation explicit, testable, and reversible, while allowing users to distinguish automatically resolved inconsistencies from records requiring expert inspection.

Applied to ITS reference data, **CurateMake** shows that CoL harmonisation resolves a substantial fraction of apparent taxonomic inconsistency, whereas SATIVA contributes additional information only when supported by taxon-coherent alignments. In this benchmark, phylogenetic curation is therefore conditional on database engineering: alignment design is not merely preprocessing, but part of the correction method itself. This conclusion is supported by two complementary evaluations: entropy-based diagnostics quantify large-scale changes in taxonomic coherence across Raw, CoL-harmonised, and SATIVA-validated annotation layers, whereas controlled error simulations isolate the behaviour of each curation layer under known error classes.

### 4.2 Interpretation of results

#### 4.2.1 A layered view of reference database curation

Reference databases are a major bottleneck in metabarcoding and metagenomic studies because both their representativeness for the ecosystem under study and their annotation quality shape downstream classification and interpretation. Evidence from metagenomic read classification shows that more comprehensive and representative databases can substantially improve classification performance (Smith et al., 2022; Méric et al., 2019; Piquer-Esteban et al., 2022), while variable database quality affects how marker-gene and whole-genome surveys are interpreted (Breitwieser et al., 2019; Pérez-Cobas et al., 2020). The same principle applies to ITS metabarcoding: reference database construction is not a neutral preprocessing step, but a component of the inference pipeline.

Reference database errors are heterogeneous. Some reflect nomenclatural drift, such as obsolete names, spelling variants, or missing ranks, whereas others reflect biological or technical inconsistencies, such as contamination, misidentification, incomplete marker recovery, or sequence-label discordance. Treating these problems as a single correction task obscures their different causes and recoverability.

The layered structure of **CurateMake** assigns these error classes to distinct but connected mechanisms. CoL harmonisation operates on names and resolves inconsistencies that can be matched to accepted taxonomy, but it cannot assess whether a sequence is biologically consistent with its label. SATIVA evaluates sequence-label consistency through phylogenetic placement, but only when the alignment and task structure provide an informative context. Preserving Raw, CoL-harmonised, and SATIVA-validated annotation layers in parallel links these mechanisms while keeping curation auditable and reversible.

This design provides a general model in which harmonisation, validation, and expert review are complementary steps rather than interchangeable alternatives (Table 3).

**Table 3:**
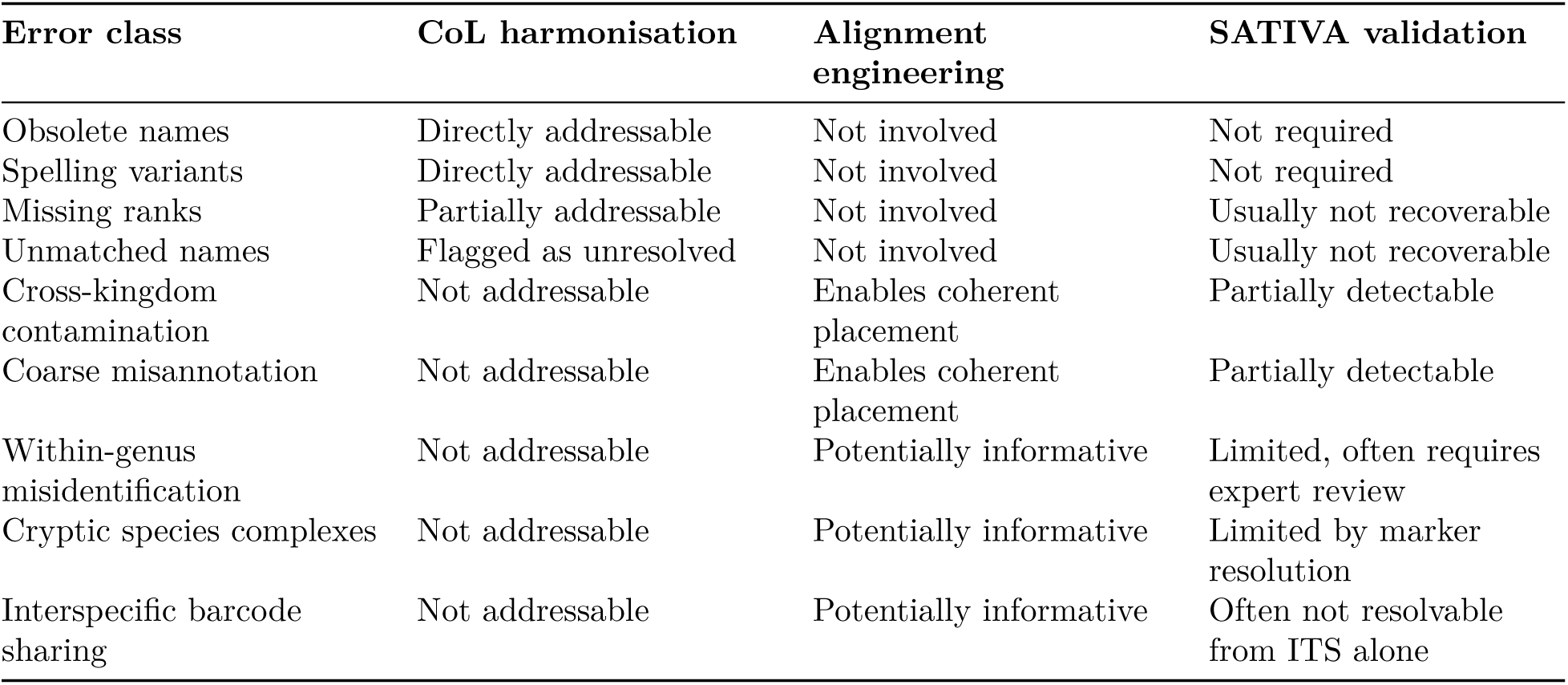
Conceptual mapping between common reference database error classes and the correction or validation layers implemented in **CurateMake**. The table describes the expected role of each layer rather than absolute performance guarantees.

#### 4.2.2 Complementarity of CoL, SATIVA, and alignment engineering

CoL and SATIVA address different parts of the taxonomic error space. In the controlled simulation, CoL harmonisation corrected 67.5% of obsolete-taxonomy errors and 42.7% of missing-rank errors at 15% corruption, confirming its value as a first-pass nomenclatural curation layer. These errors require standardised names and explicit handling of synonymies, obsolete labels, and incomplete ranks rather than phylogenetic inference.

By contrast, CoL cannot detect errors that are invisible at the level of names: a sequence may carry a valid accepted name while being contaminated, misidentified, or phylogenetically inconsistent. In the integrated workflow, SATIVA added this complementary signal, with partial contamination detection at genus or kingdom level reaching 23.4%, whereas CoL alone had no effect. The Raw, CoL-harmonised, and SATIVA-validated annotation layers should therefore be interpreted as an auditable sequence of taxonomic hypotheses, not as a single automatic truth.

A key result is that phylogeny-guided validation depends critically on alignment quality. SATIVA applied in isolation corrected 0.0% of errors across all simulation conditions, and CoL-to-SATIVA, corresponding to CoL harmonisation followed by SATIVA without the workflow’s hierarchical MSA infrastructure, matched CoL alone. SATIVA contributed only within the full workflow, where hierarchical task partitioning, taxon-level alignment, trimming, and HMM-based assignment jointly define the validation context (section 2.6). This cautions against treating SATIVA as a standalone correction module for arbitrary alignments and suggests that future gains in phylogenetic curation may come as much from better alignment engineering as from new correction algorithms.

#### 4.2.3 Taxonomic coherence and scope of inference

The entropy experiment evaluates the effect of the workflow on real datasets at scale, and, because the Raw, CoL-harmonised, and SATIVA-validated layers correspond to successive processing stages, it also tracks how label uncertainty propagates through the sequential pipeline: normalised entropy decreased monotonically from Raw to CoL to SATIVA at every rank and identity threshold, indicating that each stage reduced rather than amplified propagated uncertainty. CoL harmonisation significantly reduced intra-cluster entropy at all taxonomic ranks, indicating that much apparent label heterogeneity within sequence clusters reflects nomenclatural inconsistency. After SATIVA validation, entropy reached near-zero (*<* 10^−3^) at kingdom level, suggesting that high-level label conflicts were almost completely resolved by the full workflow. The same qualitative ordering of annotation layers was observed across the 95%, 97%, and 99% identity thresholds (figure S1), supporting the robustness of the comparison while confirming that absolute entropy values depend on clustering parameters and the composition of non-singleton clusters.

These results indicate improved taxonomic coherence, not direct taxonomic correctness. Entropy measures label consistency within sequence clusters; it does not provide ground truth or determine whether a homogeneous cluster is correctly labelled. Species-level entropy remained elevated after the complete workflow, as expected given the limits of ITS for species-level identification (Schoch et al., 2012; Kiss, 2012). Cryptic species complexes, interspecific barcode sharing, nomenclatural instability, recent taxonomic revisions, and residual database errors can all produce heterogeneity that automated harmonisation and phylogenetic placement cannot fully resolve from a single marker.

The controlled simulation was therefore essential because it evaluates known-error recovery under traceable conditions. Its results should be interpreted as relative benchmarks under defined corruption scenarios, not as estimates of natural error frequencies in production databases. The simulated dataset contains 1,989 Badread-generated reads derived from 20 ground-truth anchors and does not represent the full biological diversity of *Carex* or *Russula*. Random error injection may also be conservative for phylogeny-based methods because real database errors are often structured among related taxa, laboratory workflows, or taxonomic coverage gaps. Together, the entropy and simulation experiments answer complementary questions: Experiment 1 measures large-scale coherence in a real multi-source database, whereas Experiment 2 identifies which error classes can be recovered by each curation layer under controlled conditions.

### 4.3 Generality, reuse, and practical application

Although this study focuses on ITS, the workflow architecture is not specific to a single barcode or database source. Marker-specific components include sequence extraction, alignment strategy, and some aspects of taxonomic interpretation, whereas source normalisation, provenance tracking, nomenclatural harmonisation, phylogeny-guided validation, parallel annotation-layer retention, and coherence-based evaluation are general design principles. These components can be transferred to other barcode systems, provided that marker-specific extraction and alignment rules are adapted.

The same architecture could be adapted to COI, rbcL, matK, 16S, 18S, or other markers used in metabarcoding and environmental DNA studies. CoL could be complemented or replaced by domain-specific taxonomic authorities, and SATIVA could be replaced or supplemented by alternative phylogeny-aware validation methods. The durable contribution of **CurateMake** is therefore the auditable architecture that makes such modules interpretable, not the exclusive use of a particular taxonomic authority or validation tool (Table 4).

**Table 4:**
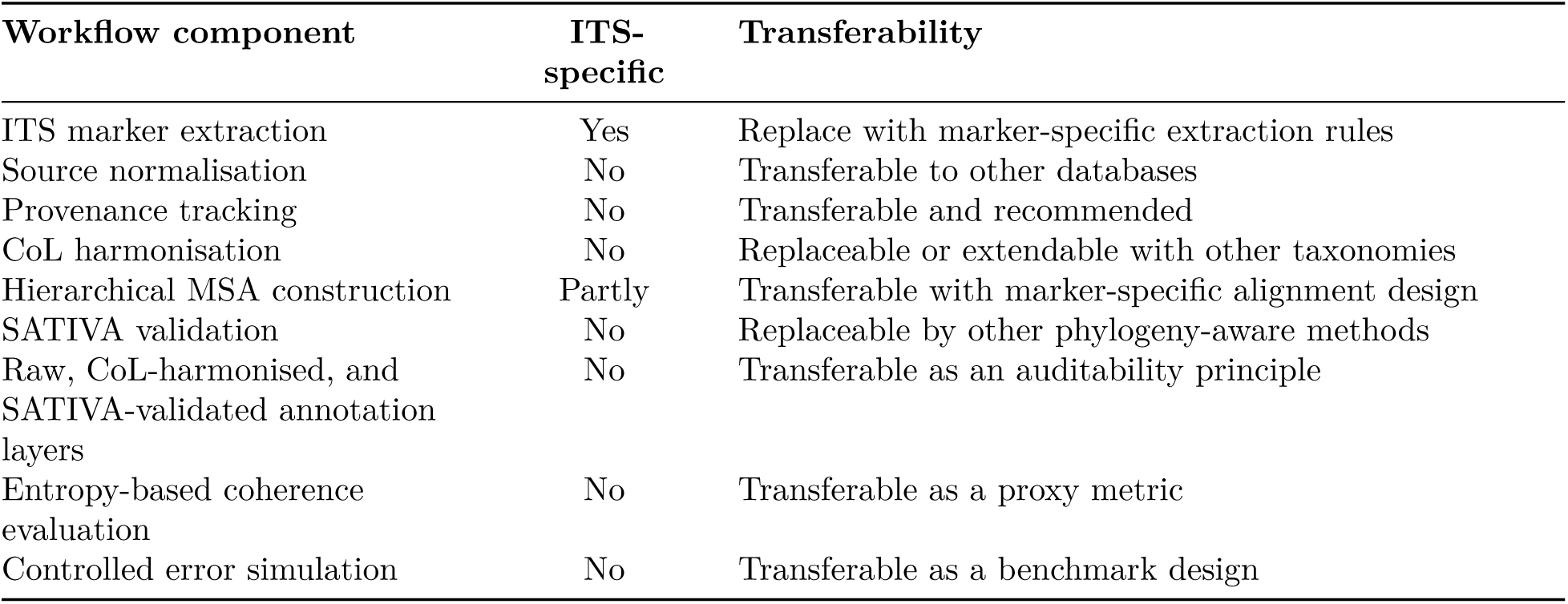
Components of **CurateMake** and their expected transferability to other metabarcoding reference database projects.

| Workflow component | ITS-specific | Transferability |
| --- | --- | --- |
| ITS marker extraction | Yes | Replace with marker-specific extraction rules |
| Source normalisation | No | Transferable to other databases |
| Provenance tracking | No | Transferable and recommended |
| CoL harmonisation | No | Replaceable or extendable with other taxonomies |
| Hierarchical MSA construction | Partly | Transferable with marker-specific alignment design |
| SATIVA validation | No | Replaceable by other phylogeny-aware methods |
| Raw, CoL-harmonised, and SATIVA-validated annotation layers | No | Transferable as an auditability principle |
| Entropy-based coherence evaluation | No | Transferable as a proxy metric |
| Controlled error simulation | No | Transferable as a benchmark design |

For ecologists and metabarcoding users, the main implication is that reference databases should not be treated as static external resources. They define the taxonomic resolution and error structure of downstream biodiversity estimates, community composition analyses, ecological indicators, and biomonitoring trends. A reproducible database construction workflow therefore makes upstream reference choices explicit and allows database versions to be compared across time.

Within this broader context, **CurateMake** is designed for projects that need to integrate multiple heterogeneous metabarcoding reference databases while preserving provenance, reproducible release history, and comparable taxonomic annotation layers. Its main use case is therefore the construction, update, and audit of reference resources assembled from sources that differ in taxonomic conventions, marker coverage, annotation completeness, or geographic scope. This is distinct from routine taxonomic assignment: the workflow structures and documents the reference database on which such assignments depend.

The current output of **CurateMake** is a structured relational database intended for audit, comparison, and downstream conversion. Developing direct exports to formats used by tools such as QIIME2, OBITools, or DADA2 would be a natural extension, allowing curated and versioned reference resources to be used more directly in metabarcoding analyses. Until such exports are implemented, **CurateMake** should be interpreted primarily as a database construction and curation workflow, not as a replacement for downstream taxonomic assignment tools or expert species-level validation.

### 4.4 Reproducibility, limitations, and outlook

**CurateMake** is implemented as a modular workflow in which source ingestion, marker extraction, nomenclatural harmonisation, MSA construction, phylogenetic validation, and database export are executed as traceable computational steps. Source provenance, Raw, CoL-harmonised, and SATIVA-validated annotation layers, software parameters, task partitioning rules, and correction outputs are retained rather than overwritten. The relational database structure and Snakemake implementation make these decisions explicit and repeatable, while supporting versioned database releases.

Several limitations remain. First, coherence is not correctness: entropy measures label consistency within sequence clusters, not absolute taxonomic accuracy. Second, controlled simulations simplify real error processes and may disadvantage phylogeny-based methods by disrupting taxon-coherent alignment structure more strongly than biologically clustered errors would. Third, species-level resolution remains limited by ITS marker resolution, cryptic diversity, interspecific barcode sharing, and unstable nomenclature. Fourth, SATIVA outputs should be interpreted as hypotheses or warnings, especially for low-abundance taxa, difficult species complexes, and cases where alignment quality or marker resolution is insufficient. Finally, full-scale execution requires substantial HPC resources: the full run required approximately two months on 190 CPUs, with ITSx and SATIVA as the dominant runtime bottlenecks. SATIVA v0.9.1 was also designed for RAxML 8.2.3, whereas all analyses here used RAxML 8.2.13; users should verify compatibility in new environments (Appendix C).

Future work should prioritise the alignment-engineering layer, because this study identifies MSA quality as a main bottleneck for phylogeny-guided validation. Alternative task partitioning strategies, alignment parameters, trimming rules, and alignment tools should be evaluated systematically, ideally with simulation outputs that record intermediate taxonomies, entropy, and error status after each processing stage. Additional priorities include incremental database updates, reuse of previous alignments, improved parallelisation, adaptation to other barcode markers, and direct export to formats used by downstream metabarcoding tools.

### 4.5 Conclusions

**CurateMake** provides a reproducible and auditable framework for harmonising and validating ITS reference databases assembled from heterogeneous sources. Rather than automating taxonomic truth, it makes reference database curation explicit, testable, and reversible by preserving Raw, CoL-harmonised, and SATIVA-validated annotation layers in parallel.

The results show that nomenclatural harmonisation and phylogeny-guided validation address different error classes: CoL resolves many name-based inconsistencies, whereas SATIVA contributes only when supported by taxon-coherent alignments. Alignment engineering is therefore a central component of phylogenetic curation, not a technical detail.

More broadly, reference database construction should be treated as part of the ecological inference pipeline. By separating taxonomic labels, clustering choices, alignment procedures, validation tools, and provenance records into auditable layers, **CurateMake** provides a practical foundation for building more consistent, reusable, and versioned reference resources for metabarcoding.

## Data Availability Statement

Raw archives are available from the original sources (BOLD, UNITE, PLANiTS, CALeDNA). Processed outputs, the curated mini-database, reproducibility scripts, and the source data underlying the supplementary tables and figures are archived on Zenodo at https://doi.org/10.5281/zenodo. 21264079.

## Code Availability Statement

**CurateMake** is available at https://github.com/Aaramis/CurateMake under the GNU General Public License v3.0. The version used in this manuscript is release v1.0.0 (commit 603b5a2), archived on Zenodo at https://doi.org/10.5281/zenodo.21263905.

## Author Contributions

- Auguste Gardette: Conceptualization, Methodology, Software, Formal Analysis, Writing – Original Draft.
- Eugeni Belda: Validation, Data Curation, Writing – Review & Editing.
- Edi Prifti: Writing – Review & Editing, Supervision.
- Jean-Daniel Zucker: Supervision, Funding Acquisition.

## Acknowledgements

We thank the IFB (Institut Français de Bioinformatique) clusters for computational resources.

## Conflict of Interest

The authors declare no conflict of interest.

**.1 Justification: potential as a benchmark dataset**

Provided for editors and reviewers, per the Methods in Ecology and Evolution Workflow Checklist.

We argue that the datasets released with **CurateMake** could serve as reusable community benchmarks for reference-database curation, for five reasons.

**(1) They fill a documented gap.** No standard benchmark currently exists for evaluating multi-source ITS reference-database harmonisation and curation: existing resources are either single-source repositories or read-processing test sets (Table 1). The lack of a shared benchmark is itself a barrier to comparing curation methods.
**(2) Scale and cross-kingdom coverage.** The full dataset integrates four major public repositories (UNITE, BOLD, PLANiTS, CALeDNA) into 3,581,943 ingested records, expanded to 5,189,645 barcode-resolved entries spanning 7 kingdoms and ≈183,000 species-level labels. This breadth exceeds what any single source provides and exposes the heterogeneous, cross-kingdom label conflicts that a curation benchmark must contain.
**(3) Structured provenance makes it evaluable, not merely large.** Every record retains links to its source and three parallel annotation layers (Raw, CoL-harmonised, SATIVA-validated). A new curation or classification method can therefore be scored against three defined reference states and against record-level provenance, which a raw sequence dump does not allow.
**(4) A companion ground-truth benchmark.** The controlled error-injection dataset (20 *Carex*/*Russula* anchors, 1,989 reads, five error types, eight corruption levels × five seeds) provides fully traceable known-error scenarios and ready-made evaluation protocols (correction rate by error type; normalised intra-cluster entropy). Both metrics are directly reusable to score other methods.
**(5) Open, versioned, and regenerable.** The data are archived under an open DOI (https://doi.org/10.5281/zenodo.21264079) and the GPLv3 workflow regenerates, extends, and versions them (new sources, updated releases, other markers), the prerequisites for a maintainable “living” benchmark rather than a one-off snapshot.

We note explicitly that the value of these datasets as a benchmark rests on traceable provenance, reproducibility, and the simulated ground truth, not on a claim of absolute taxonomic gold-standard truth: as discussed, intra-cluster coherence is a proxy for, not a proof of, taxonomic correctness.

## A Reproducibility metadata

This appendix summarises the dataset versions, sequence counts, software versions, and key parameters used in the workflow. Dataset counts correspond to records retained after ingestion and pipeline-level filtering, rather than to raw download sizes. Software versions and parameters were extracted from the workflow configuration files, Conda environments, runtime modules, and pipeline scripts.

### A.1 Data sources and dataset composition

All public reference datasets were extracted in late August 2025, although public release dates differed among sources (main-text Table 2). The full dataset was used for large-scale entropy analyses, whereas the public mini-dataset was used for demonstration, testing, and runtime benchmarking. Detailed barcode-level counts are provided in Table S1.

**Table S1:**
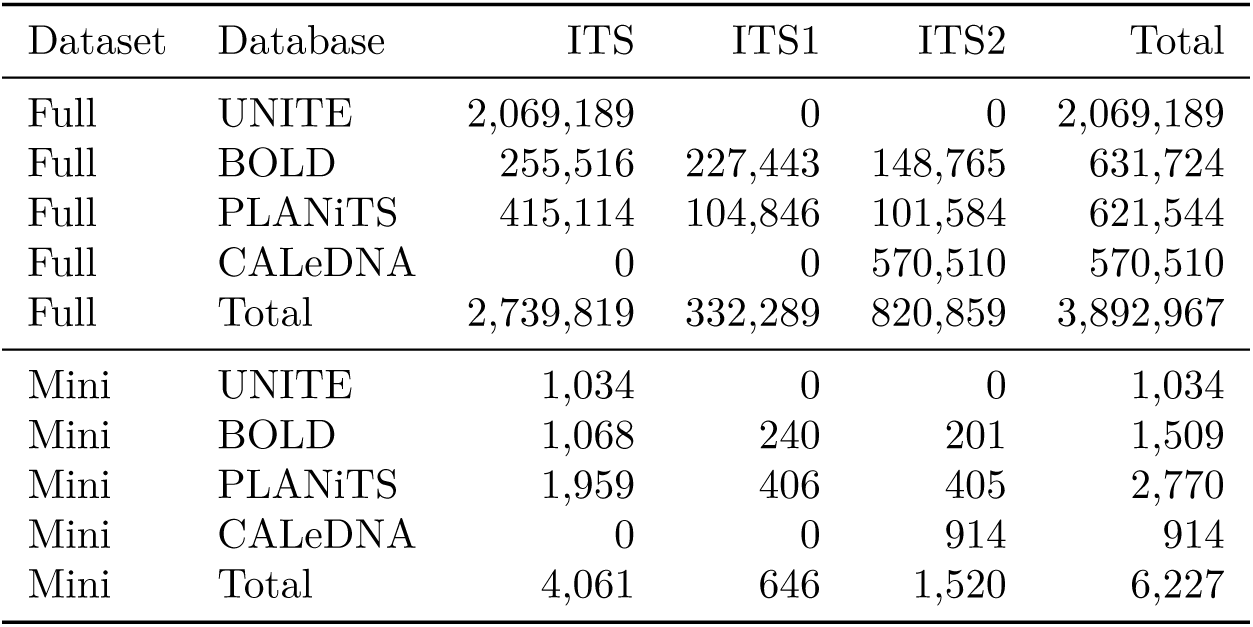
Barcode-level record counts after ingestion and pipeline filtering for the full and mini datasets. These counts are stratified by parsed barcode type and are not source-record totals: a source record can contribute more than one barcode-level record when multiple barcode annotations are retained.

### A.2 Controlled error-injection simulation (Experiment 2)

This section provides the reproducibility details of the controlled error-injection simulation summarised in the Methods and Results. The benchmark was designed to compare correction strategies under fully traceable ground truth rather than to characterise taxon-wide ecological performance.

#### A.2.1 Ground-truth anchor sequences

Twenty full-ITS sequences were manually retrieved from BOLD in January 2026 and used as ground-truth anchors: 10 plant species in *Carex* (Cyperaceae) and 10 fungal species in *Russula* (Russulaceae) (Table S2). These genera provide complementary test contexts for cross-kingdom contamination and within-genus misidentification. Species selection was restricted to entries with a CoL-accepted taxonomy and an unambiguous BOLD record, so that recovery could be evaluated against verifiable ground truth rather than confounded by nomenclatural ambiguity.

**Table S2:**
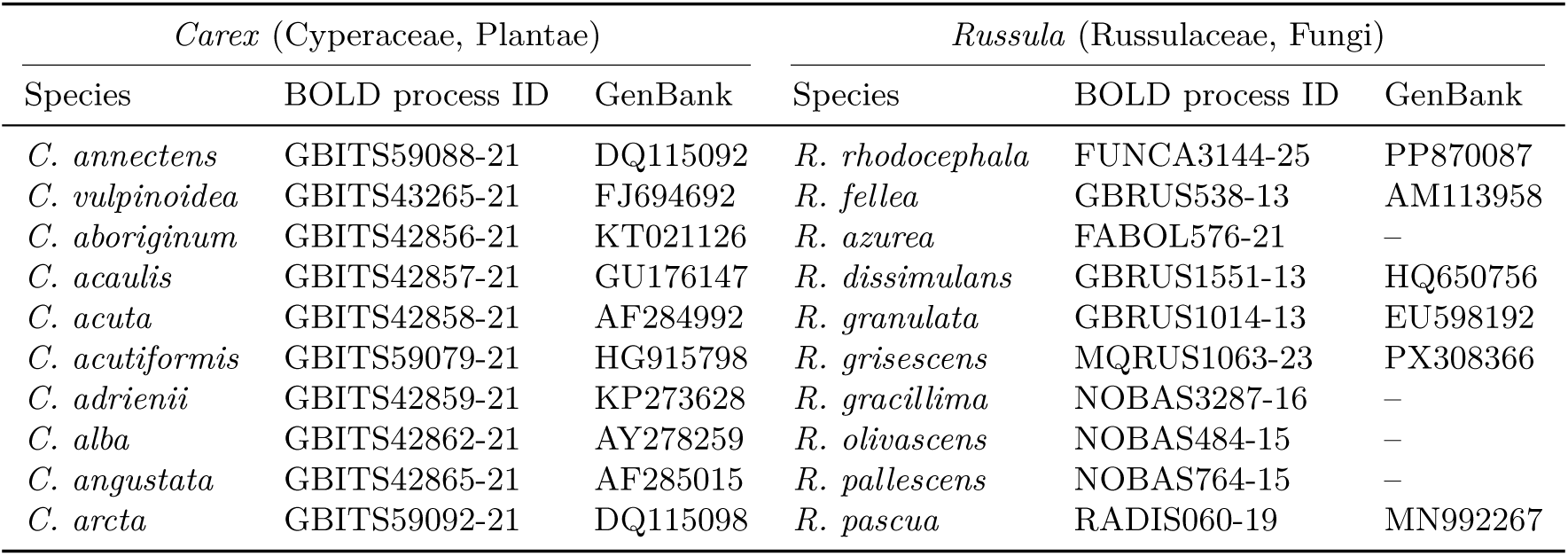
Ground-truth anchor species used to build the controlled error-injection simulation. All species were confirmed as CoL-accepted at retrieval. **BOLD process ID** identifies the retrieved full-ITS record on BOLD; **GenBank** gives the corresponding INSDC nucleotide accession where available (a dash indicates records without a linked INSDC accession).

#### A.2.2 Sequence augmentation

Each of the 20 anchor sequences was amplified *in silico* with Badread (Wick, 2019) using –identity 97,99,2, that is, a target mean read identity of 97%, a maximum of 99%, and a standard deviation of 2%. Read length was fixed to the anchor sequence length (length standard deviation set to zero), and no junk reads or chimeras were generated. A total of 201 reads were simulated per species. After retaining only positive-strand reads to mimic amplicon directionality, the augmented dataset contained 1,989 sequences. Corruptions were applied only to the taxonomy fields; the DNA sequences themselves were never modified.

#### A.2.3 Taxonomy-corruption types

For each scenario, a configurable proportion of sequences was sampled at random and assigned one of five error types with uniform probability by default. Corruption rates were 1%, 2%, 3%, 5%, 10%, 15%, 25%, and 50%, each run with five independent seeds, yielding 40 scenarios. All corruptions were recorded in per-sequence reports storing the original and corrupted taxonomies. The five error types are defined as follows:

##### MISSING_RANK

One to three ranks, excluding kingdom, are replaced by Na.

##### OBSOLETE_TAXONOMY

One rank is replaced by a random character string, emulating a deprecated or non-standard name.

##### MISIDENTIFICATION

The species label is replaced by that of another species within the same genus.

##### CONTAMINATION

The full taxonomy is replaced by that of another organism, drawn from a different kingdom when possible.

##### UNIDENTIFIABLE

All ranks from a randomly chosen level (order through species) downward are replaced by Na.

#### A.2.4 Correction strategies (component ablation)

Because taxonomic annotations are modified only by CoL harmonisation and SATIVA integration, each corrupted dataset was processed under four strategies that isolate the contributions of nomenclatural harmonisation, phylogenetic validation, and the workflow’s alignment-engineering block:

##### Full pipeline

CoL harmonisation, ITSx extraction, hierarchical MSA/HMM-based alignment assignment, and SATIVA validation — the complete **CurateMake** workflow.

##### CoL alone

Catalogue of Life harmonisation only, with no phylogenetic validation.

##### SATIVA alone

Kingdom-level MAFFT alignments and SATIVA, without prior CoL harmonisation and without the workflow’s hierarchical HMM-based alignment assignment.

##### CoL-to-SATIVA

CoL harmonisation followed by SATIVA, but without the workflow’s hierarchical HMM-based alignment assignment.

Per-error-type correction was defined as the proportion of corrupted records of a given type fully restored to ground-truth taxonomy (all corrupted ranks corrected). For `CONTAMINATION`, where full-rank correction was negligible, partial detection was reported instead as the mean of genus-level and kingdom-level recovery. The global (overall) correction rate is the unweighted mean of the four per-error-type rates (`OBSOLETE_TAXONOMY, MISSING_RANK, CONTAMINATION` as partial detection, and `MISIDENTIFICATION`), computed per seed and then averaged across seeds. `UNIDENTIFIABLE` errors yield null correction rates by construction and are excluded from both the per-error-type panel and the overall rate. Means and standard errors were computed across the five seeds for each corruption rate.

#### A.2.5 Correction results

Tables S3 and S4 report the numerical values underlying figure 3. All values are means ± SEM over the five matched seeds (seeds 1–5, for which all four strategies were run). The overall correction rate is the unweighted mean of the four per-error-type rates (`OBSOLETE_TAXONOMY, MISSING_RANK, CONTAMINATION` as partial detection, `MISIDENTIFICATION`), computed per seed before averaging. Per-seed values are available in the Zenodo deposit (`simulations_summary.csv`).

**Table S3:**
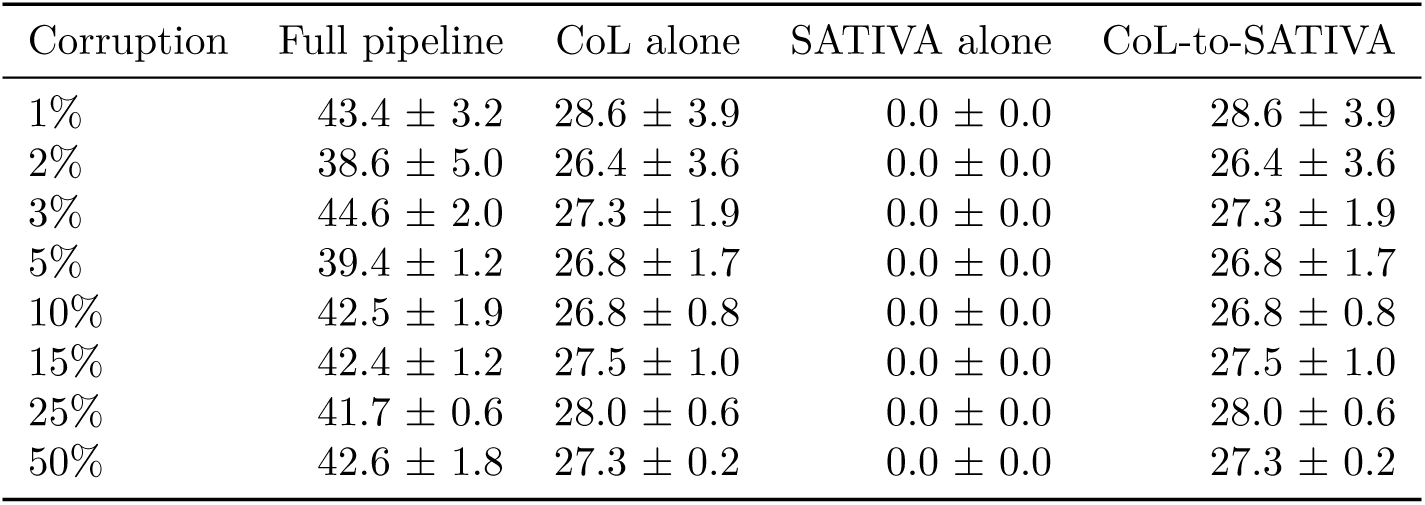
Overall correction rate (%) by taxonomic corruption rate for the four correction strategies (mean ± SEM, *n* = 5 seeds), corresponding to figure 3A. **CurateMake** (full pipeline) is the highest-performing strategy at every corruption level. SATIVA alone corrects nothing, and CoL-to-SATIVA is identical to CoL alone, confirming that phylogenetic validation contributes only within the workflow’s alignment infrastructure.

| Corruption | Full pipeline | CoL alone | SATIVA alone | CoL-to-SATIVA |
| --- | --- | --- | --- | --- |
| 1% | 43.4 ± 3.2 | 28.6 ± 3.9 | 0.0 ± 0.0 | 28.6 ± 3.9 |
| 2% | 38.6 ± 5.0 | 26.4 ± 3.6 | 0.0 ± 0.0 | 26.4 ± 3.6 |
| 3% | 44.6 ± 2.0 | 27.3 ± 1.9 | 0.0 ± 0.0 | 27.3 ± 1.9 |
| 5% | 39.4 ± 1.2 | 26.8 ± 1.7 | 0.0 ± 0.0 | 26.8 ± 1.7 |
| 10% | 42.5 ± 1.9 | 26.8 ± 0.8 | 0.0 ± 0.0 | 26.8 ± 0.8 |
| 15% | 42.4 ± 1.2 | 27.5 ± 1.0 | 0.0 ± 0.0 | 27.5 ± 1.0 |
| 25% | 41.7 ± 0.6 | 28.0 ± 0.6 | 0.0 ± 0.0 | 28.0 ± 0.6 |
| 50% | 42.6 ± 1.8 | 27.3 ± 0.2 | 0.0 ± 0.0 | 27.3 ± 0.2 |

**Table S4:**
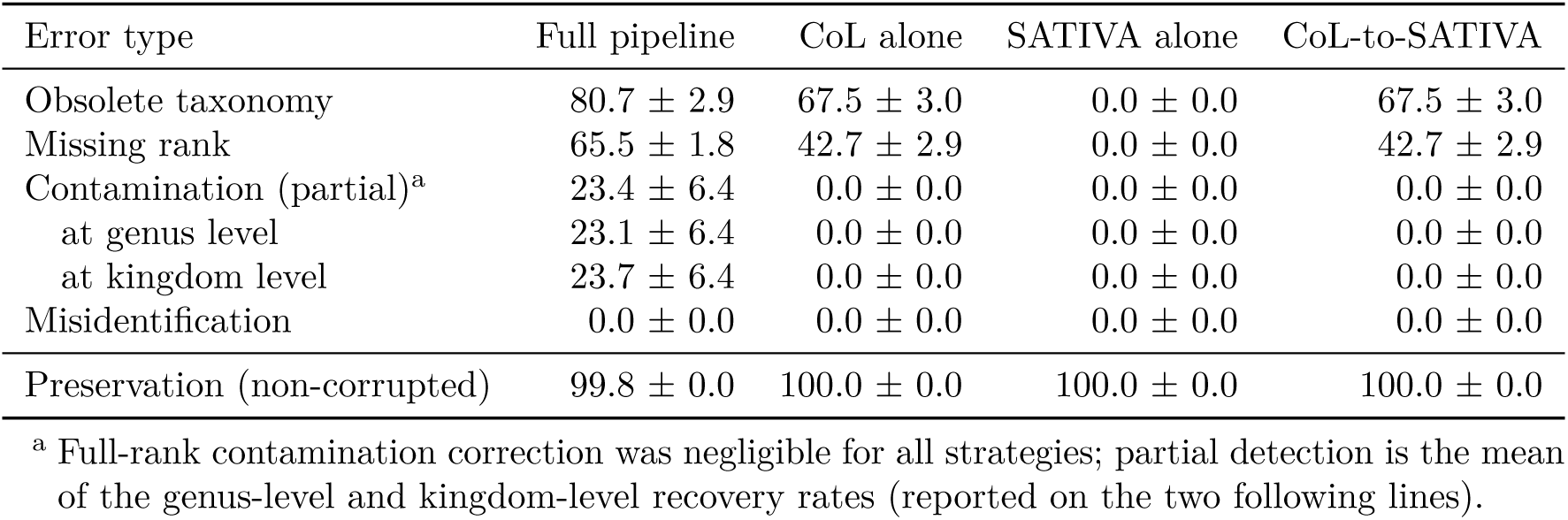
Per-error-type correction rate (%) at 15% corruption for the four strategies (mean ± SEM, *n* = 5 seeds), corresponding to figure 3B, together with the preservation rate of non-corrupted records. UNIDENTIFIABLE errors are omitted (null correction by construction). CoL-to-SATIVA reproduces CoL alone exactly.

| Error type | Full pipeline | CoL alone | SATIVA alone | CoL-to-SATIVA |
| --- | --- | --- | --- | --- |
| Obsolete taxonomy | 80.7 ± 2.9 | 67.5 ± 3.0 | 0.0 ± 0.0 | 67.5 ± 3.0 |
| Missing rank | 65.5 ± 1.8 | 42.7 ± 2.9 | 0.0 ± 0.0 | 42.7 ± 2.9 |
| Contamination (partial)<sup>a</sup> | 23.4 ± 6.4 | 0.0 ± 0.0 | 0.0 ± 0.0 | 0.0 ± 0.0 |
| at genus level | 23.1 ± 6.4 | 0.0 ± 0.0 | 0.0 ± 0.0 | 0.0 ± 0.0 |
| at kingdom level | 23.7 ± 6.4 | 0.0 ± 0.0 | 0.0 ± 0.0 | 0.0 ± 0.0 |
| Misidentification | 0.0 ± 0.0 | 0.0 ± 0.0 | 0.0 ± 0.0 | 0.0 ± 0.0 |
| Preservation (non-corrupted) | 99.8 ± 0.0 | 100.0 ± 0.0 | 100.0 ± 0.0 | 100.0 ± 0.0 |

<sup>a</sup> Full-rank contamination correction was negligible for all strategies; partial detection is the mean of the genus-level and kingdom-level recovery rates (reported on the two following lines).

**Table S5:**
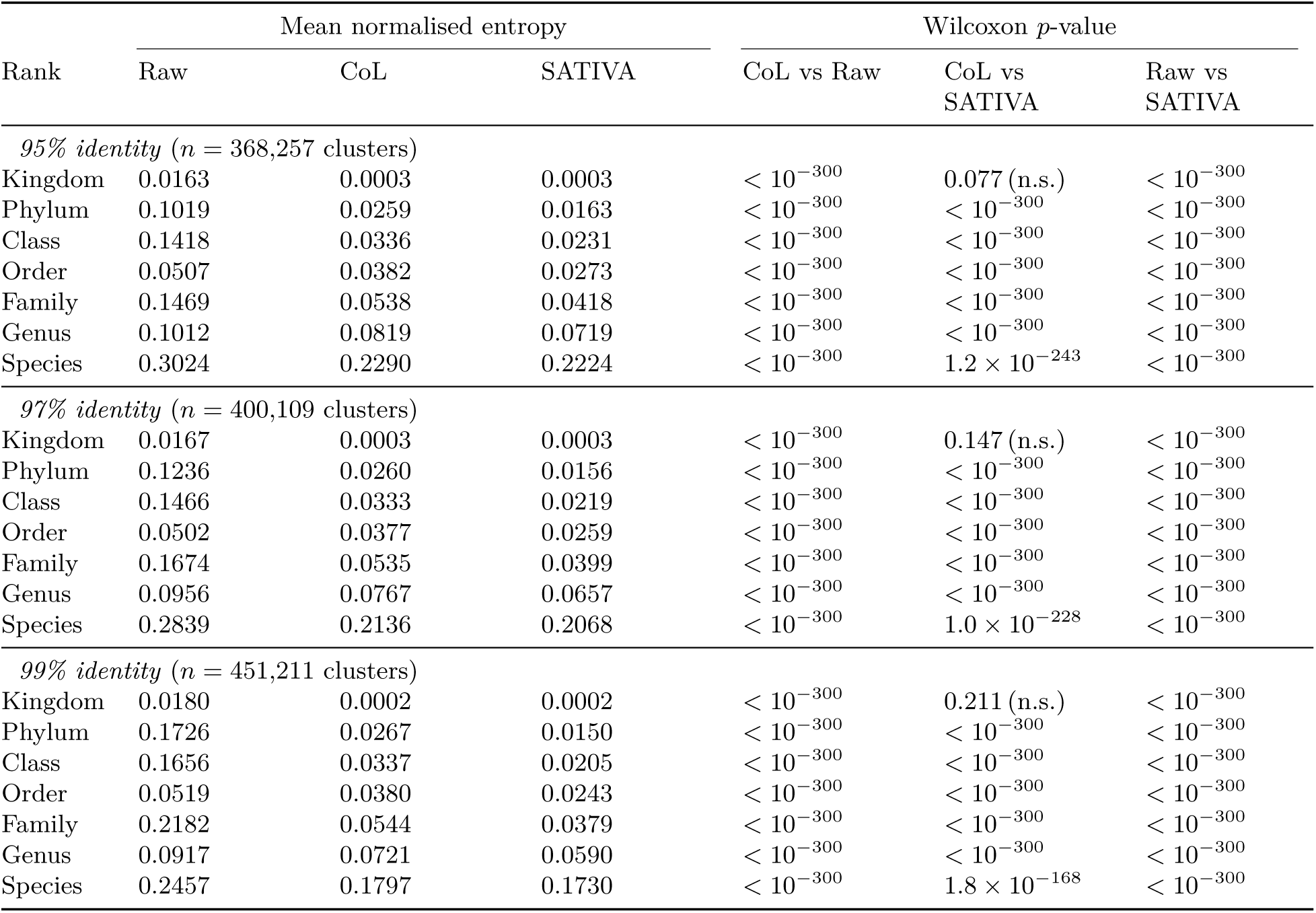
Complete intra-cluster entropy comparison across annotation layers (Experiment 1). For each clustering identity threshold (95%, 97%, 99%) and taxonomic rank, columns **Raw**, **CoL**, and **SATIVA** give the mean normalised intra-cluster entropy (*H/* log_2_ *N*) across matched non-singleton clusters. The three rightmost columns give the Wilcoxon signed-rank test *p*-values for the pairwise layer comparisons (CoL vs Raw, CoL vs SATIVA, Raw vs SATIVA), computed on matched cluster entropies. *n* is the number of matched non-singleton clusters at each threshold; this count is rank-independent because missing labels are retained as an “Unknown” category rather than excluded (see Methods). Values reported as *<* 10^−300^ fell below double-precision floating-point resolution; “n.s.” denotes non-significant comparisons (*p* ≥ 0.05). Entropy decreases monotonically from Raw to CoL to SATIVA at every rank and threshold, and the only non-significant comparison is CoL vs SATIVA at kingdom level, consistent with CoL alone resolving most kingdom-level conflicts (see figure 2).

**Figure S1:**
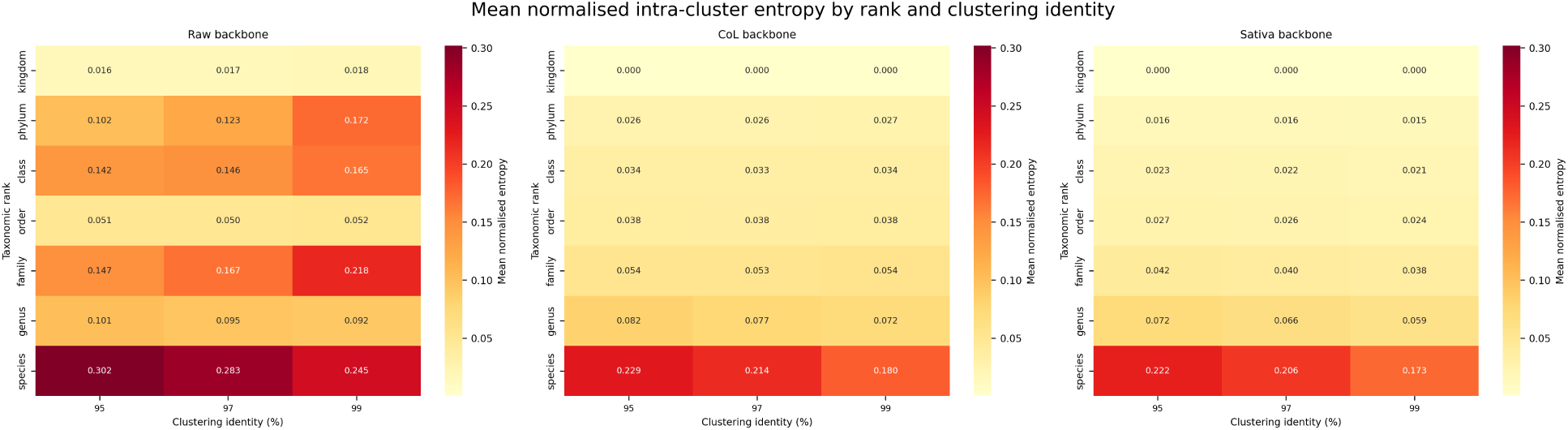
Mean normalised intra-cluster entropy (*H/* log_2_ *N*) per taxonomic rank and clustering identity threshold (95%, 97%, 99%), for each taxonomic annotation layer (Raw, CoL, SATIVA). Cell values are mean normalised entropy across all eligible non-singleton clusters. The species rank shows the highest residual entropy across all annotation layers and thresholds. The relative ordering of annotation layers (Raw *>* CoL *>* SATIVA) and the rank-entropy profile shape are qualitatively preserved across the three identity thresholds, supporting the robustness of the comparison presented in figure 2.

**Table S6:**
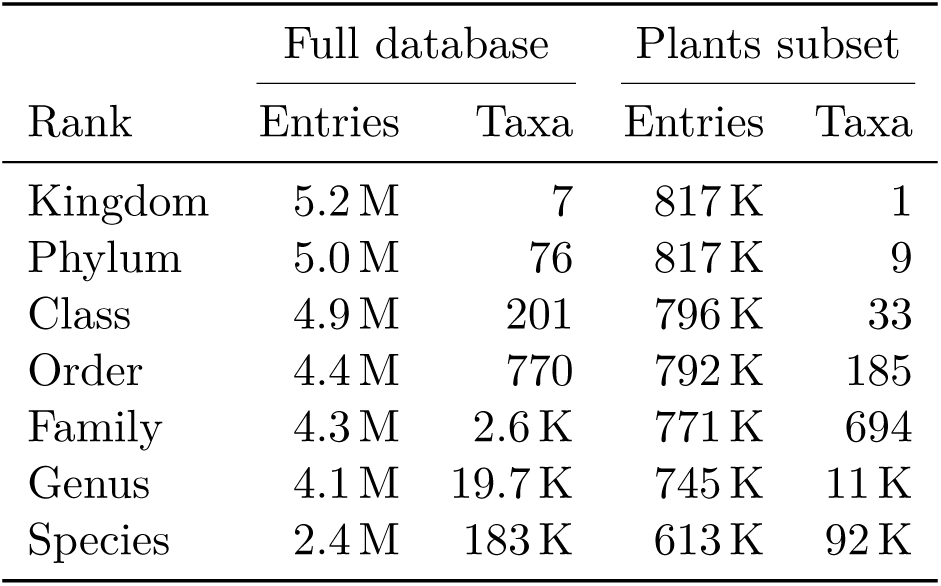
Taxonomic composition of the final harmonised database. For each rank, **Entries** is the number of barcode-resolved entries carrying a non-empty annotation at that rank, and **Taxa** is the number of distinct taxon names observed at that rank. Counts use the consolidated annotation per entry (SATIVA suggestion where available, otherwise CoL, otherwise Raw). The *Full database* columns cover all 5.19 M barcode-resolved entries (one entry per ITS/ITS1/ITS2 sub-region, so a full-length ITS sequence contributes several entries); the *Plants subset* columns are restricted to entries whose consolidated kingdom is Plantae. Entry counts decrease towards finer ranks as annotation becomes incomplete, while the number of distinct taxa increases. The species rank is the least complete: 2.4 M of 5.2 M entries (≈46%) in the full database and 613 K of 817 K (≈75%) in the plants subset are resolved to species.

**Figure S2:**
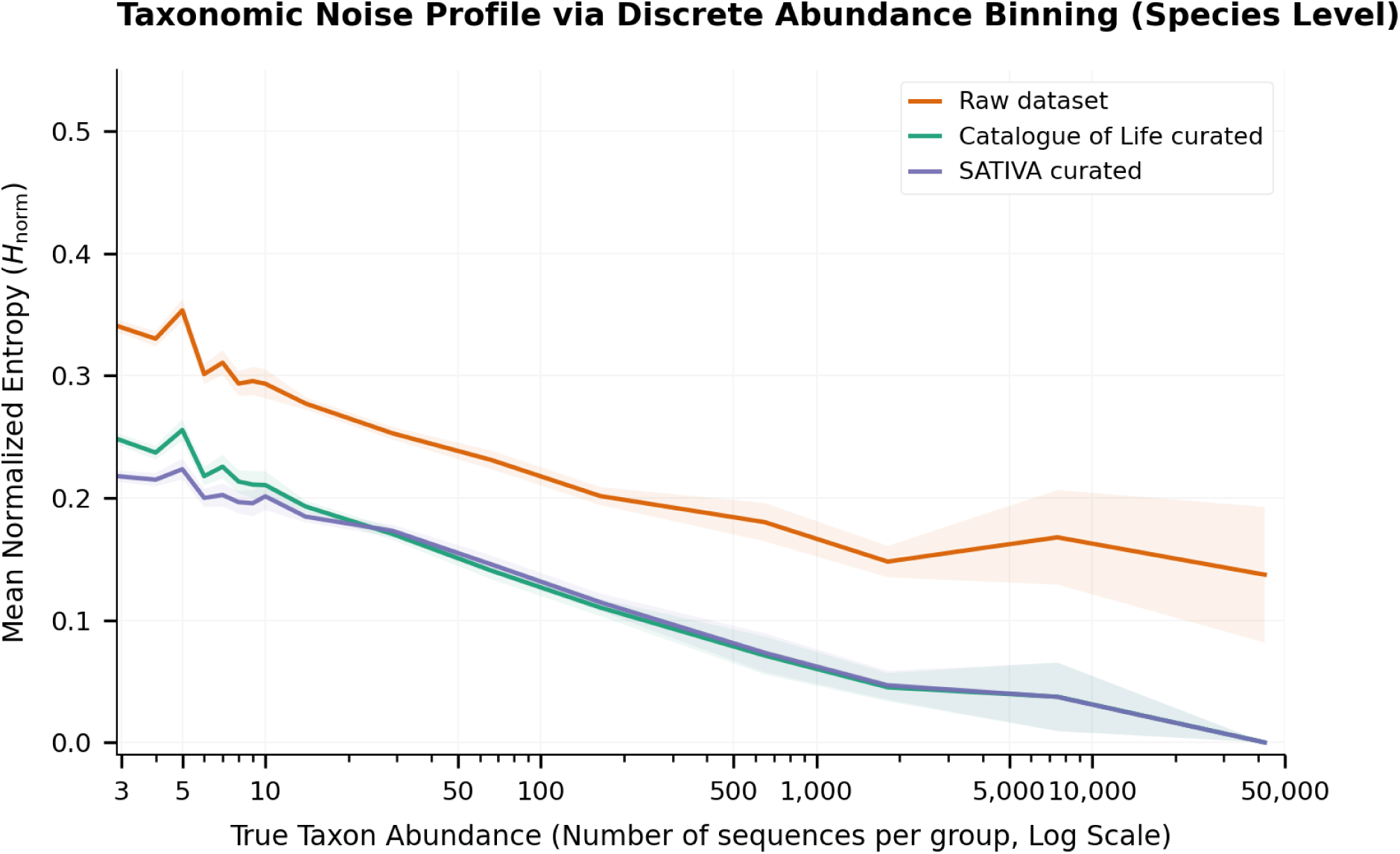
Mean normalised species-level intra-cluster entropy (*H/* log_2_ *N*) as a function of true taxon abundance (number of sequences per taxon, log scale), for each taxonomic annotation layer (Raw, CoL, SATIVA). For all annotation layers, residual entropy decreases with taxon abundance: abundant taxa are curated to near-zero entropy, whereas rare taxa retain substantial heterogeneity. Most of the reduction relative to Raw is contributed by CoL harmonisation. SATIVA’s additional reduction beyond CoL — the gap between the CoL and SATIVA curves — is concentrated at low abundance, where CoL leaves residual entropy; for abundant taxa, CoL already drives entropy close to zero, leaving little margin for further correction. This pattern is consistent with a floor effect (CoL saturating the achievable reduction in well-represented taxa) rather than a loss of phylogenetic signal in large groups. Entropy is measured at the species level, whereas MSA tasks are built at coarser ranks (family/genus, ≥ 20 sequences): a rare species can therefore still receive a SATIVA species-level correction within a larger family- or genus-level alignment.

**Table S7:**
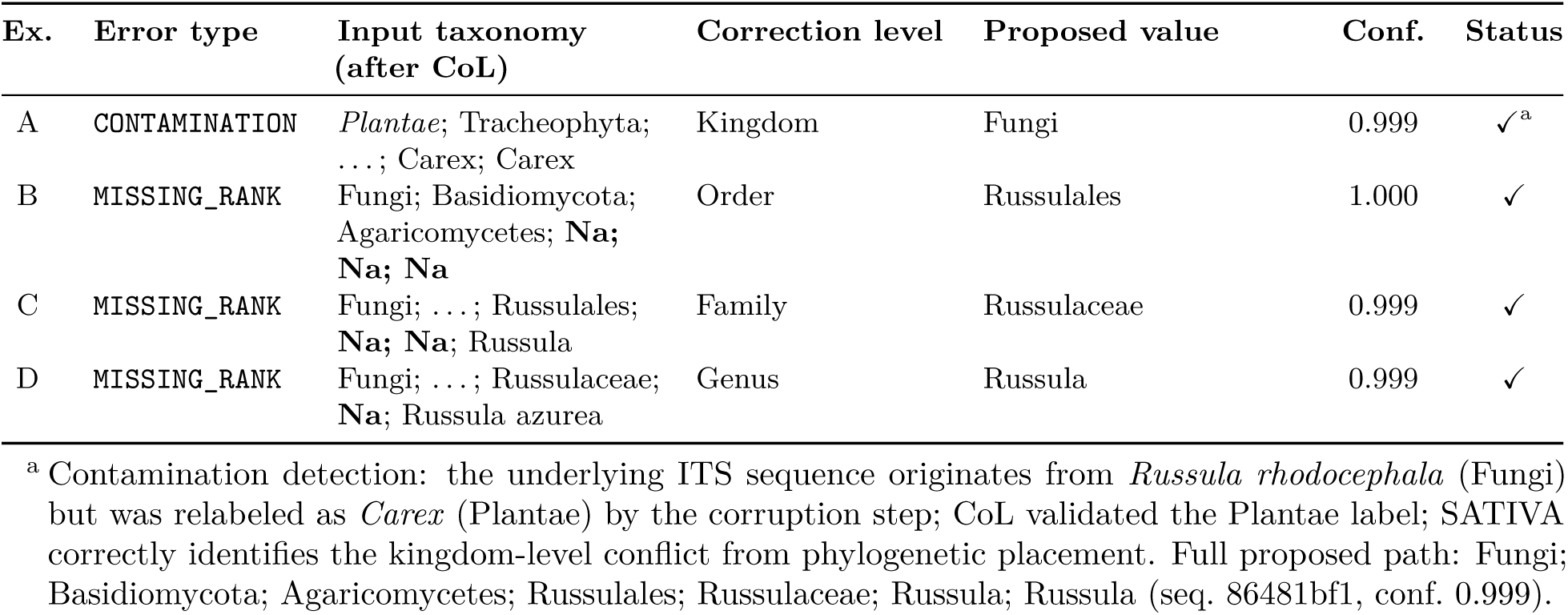
Representative SATIVA correction examples from the controlled error simulation (seed 1, 15% corruption rate), under the full pipeline. Each row shows a sequence that was corrupted during the simulation, processed through CoL harmonisation, and then flagged and corrected by SATIVA. **Input taxonomy** is the label assigned after CoL harmonisation (before SATIVA); intermediate lineage ranks omitted with ‘. . . ’ for clarity. **Correction level** is the highest rank at which SATIVA identified a mislabeling. **Proposed value** is SATIVA’s suggestion for the mislabeled rank and all ranks below it. **Status**: ✓ = full agreement with ground-truth at that rank. All four examples are from the Carex/Russula simulation dataset; the underlying ITS sequences are Badread-generated synthetic reads from 20 ground-truth anchors. Seq. IDs are truncated to the first 8 characters.

**Table S8:**
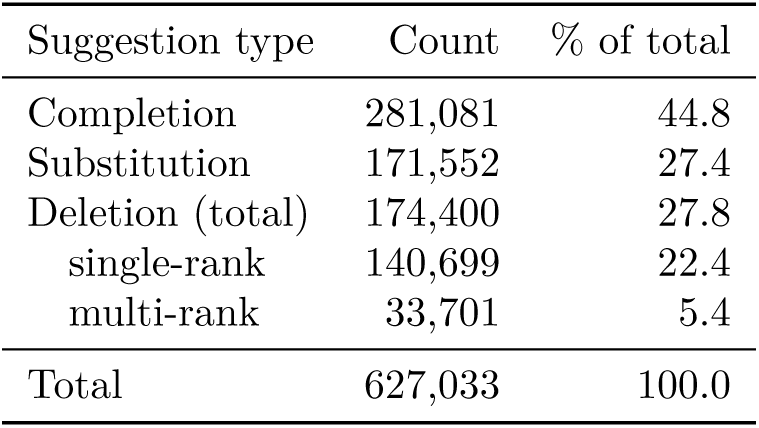
Distribution of SATIVA taxonomic suggestions on the full dataset, by suggestion type (`sativa_warnings.sativa_modification_type`). SATIVA produced 627,033 suggestions in total. **Completions** add one or more previously empty ranks below the validated lineage; **substitutions** replace an existing label at the deepest shared rank; **deletions** remove annotation and are split into single-rank (`DELETION_SIMPLE`) and multi-rank (`DELETION_MULTIPLE`) suggestions. Completions and substitutions are the most directly actionable classes; deletions are treated more cautiously because they remove existing annotation.

**Table S9:**
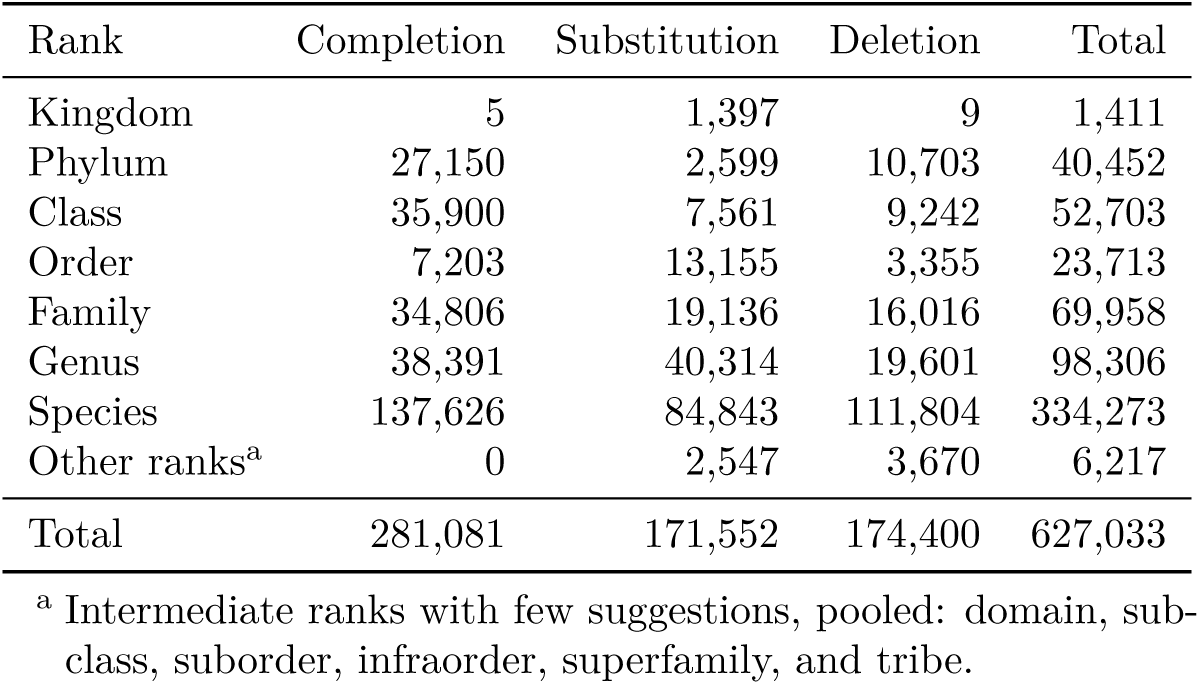
SATIVA suggestions on the full dataset by taxonomic rank of the correction (`sativa_warnings.mislabeled_level`) and suggestion type. Species is by far the most frequently corrected rank (334,273 suggestions, 53% of the total), consistent with SATIVA operating mostly on fine-rank labels. Completions dominate at intermediate ranks (phylum–family), whereas substitutions and deletions are concentrated at genus and species level.

**Table S10:**
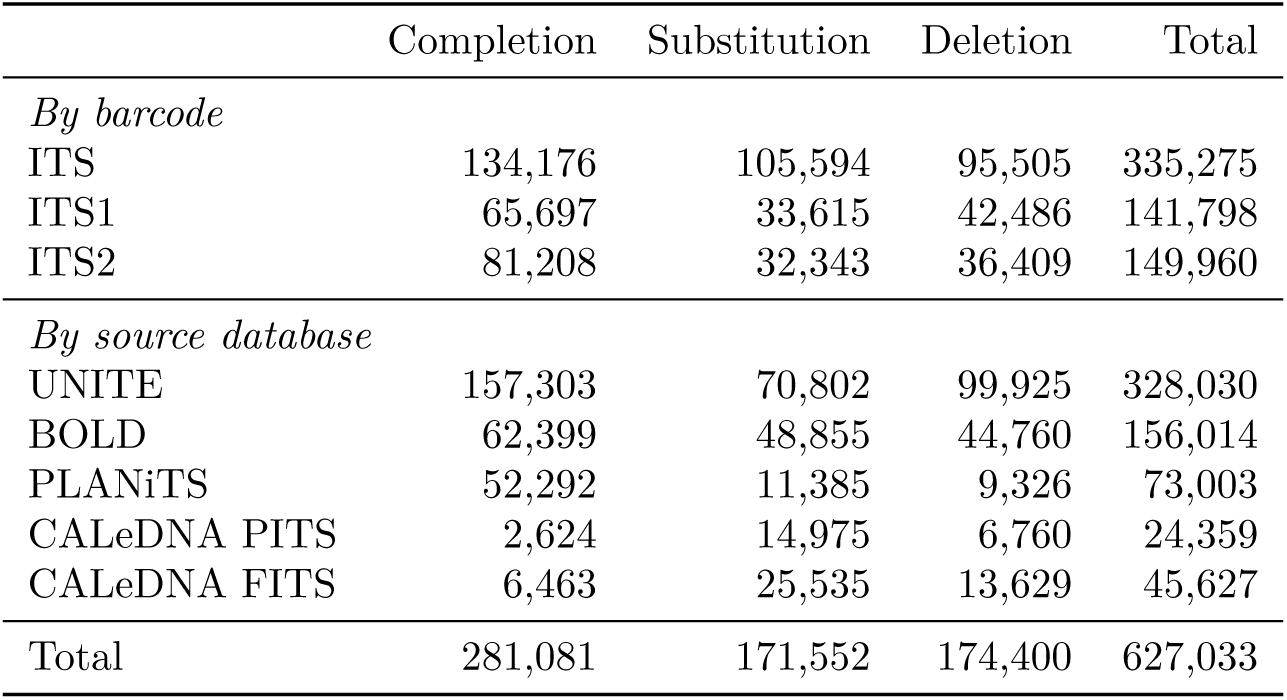
SATIVA suggestions on the full dataset by barcode type and by source database, split by suggestion type. Counts are numbers of suggestions (`sativa_warnings`); deletions pool `DELETION_SIMPLE` and `DELETION_MULTIPLE`. Suggestions arise across all barcodes and all sources. UNITE, the largest source, contributes the most suggestions, and completions are the leading type for UNITE, BOLD, and PLANiTS, whereas the CALeDNA subsets — which enter with coarse “Eukaryota” kingdom labels — yield proportionally more substitutions.

**Table S11:**
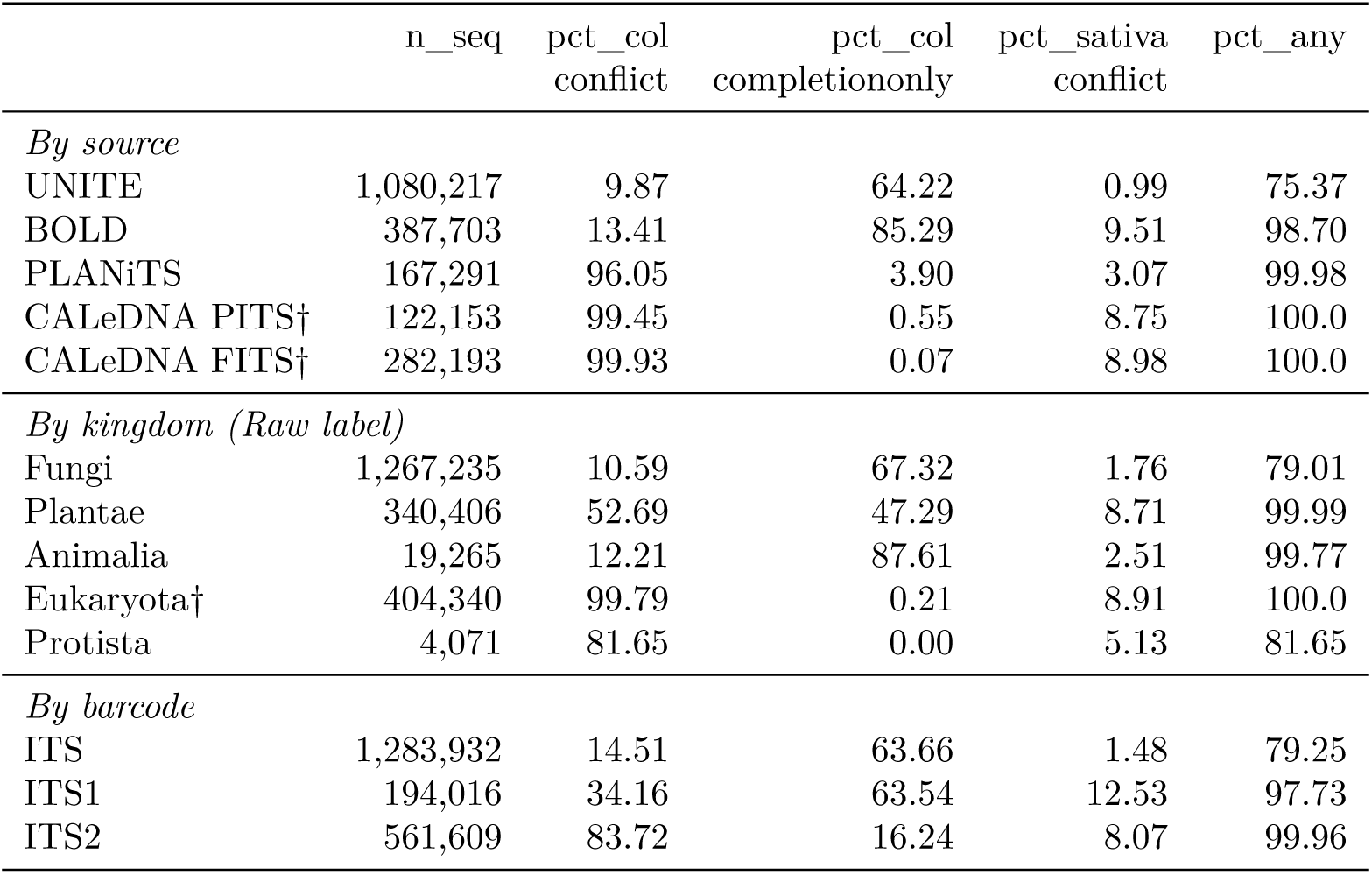
Taxonomic modification rates per source, kingdom, and barcode for original sequences (parent_seq_id IS NULL, excluding duplicates). **pct_col_conflict**: proportion of sequences where CoL replaced at least one non-empty Raw rank with a different non-empty value (nomenclatural conflict). **pct_col_completiononly**: CoL changed the lineage but only by filling previously empty ranks (no conflict). **pct_sativa_conflict**: SATIVA replaced at least one non-empty CoL rank with a different non-empty value (phylogenetic re-assignment). **pct_sativa_mod**: any SATIVA modification (including rank completions by SATIVA). **pct_any**: final taxonomy (Sativa *>* CoL *>* Raw) differs from Raw at any rank. All rates are conservative lower bounds (pipeline-detected changes only); a conflict reflects a detected nomenclatural difference, not necessarily an annotation error. † CALeDNA systematically labels sequences as kingdom “Eukaryota” rather than the CoL-accepted “Fungi”/“Plantae”; the near-100% conflict rate reflects this labelling convention difference, not random misannotation.

**Table S12:**
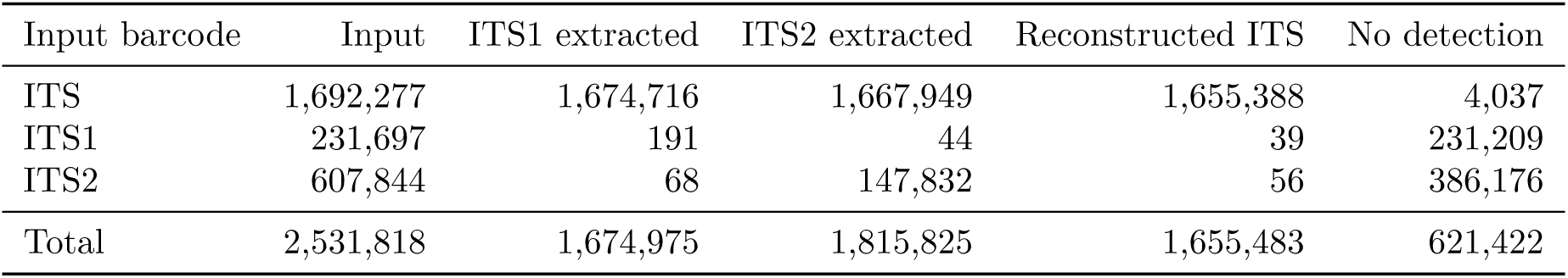
ITSx extraction on the full dataset, by input barcode type (numerical values underlying the centre panel of figure 4). **Input** is the number of deduplicated sequences of each barcode type entering ITSx; **ITS1** and **ITS2 extracted** are the sub-regions detected and re-inserted; **Reconstructed ITS** counts full ITS regions rebuilt from co-extracted ITS1/ITS2 sharing a parent; **No detection** is the number of sequences in which ITSx found no region (these are retained as pass-through records). Almost all extraction comes from full-length ITS inputs; pre-trimmed ITS1/ITS2 inputs rarely yield sub-regions because they lack the conserved flanking regions ITSx relies on.

**Table S13:**
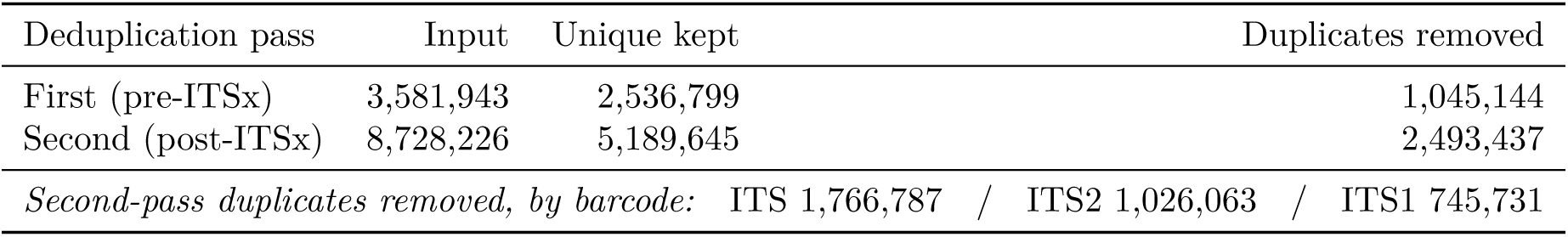
Deduplication summary on the full dataset (100% identity over full-length sequences with identical taxonomy). The first pass, before ITSx, removes source-level redundancy; the second pass, after ITSx, removes duplicates introduced by sub-region extraction and reconstruction. **Duplicates removed by barcode** shows how the removed sequences of the second pass split across barcode types. The final 5,189,645 unique barcode-resolved entries constitute the harmonised database.

**Table S14:**
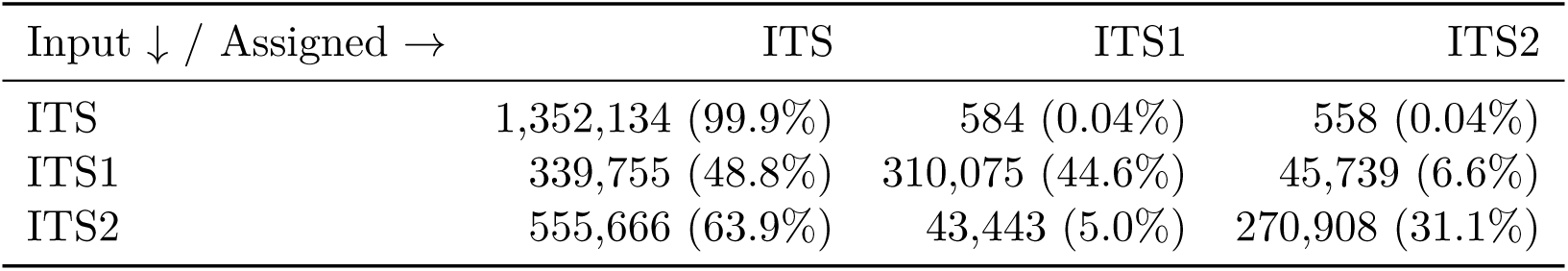
HMM-based barcode assignment on the full dataset (hmmscan, 2,918,862 sequences). Rows are the input barcode label at ingestion; columns are the barcode type assigned by the HMM library for alignment construction. Percentages are row-normalised (proportion of each input barcode type assigned to each output type). This step organises sequences with their close neighbours for alignment; it does not modify taxonomy. The high ITS1/ITS2 → ITS rates reflect inclusive matching of full-ITS profiles to sub-region queries (ITS1 and ITS2 are nested within the full ITS), not recovery of full-length signal; downstream analyses retain the original ingestion barcode.

| Input ↓ / Assigned → | ITS | ITS1 | ITS2 |
| --- | --- | --- | --- |
| ITS | 1,352,134 (99.9%) | 584 (0.04%) | 558 (0.04%) |
| ITS1 | 339,755 (48.8%) | 310,075 (44.6%) | 45,739 (6.6%) |
| ITS2 | 555,666 (63.9%) | 43,443 (5.0%) | 270,908 (31.1%) |

## B Supplementary Figures

## C Software versions and parameters

This annex summarises tool versions (as configured) and the exact parameters used in the pipeline. Values are extracted from code and configuration files.

### C.1 A) Software versions (as configured)

#### Sources

- Conda environment files: workflow/envs/*.yaml
- SATIVA bundle: workflow/scripts/Sativa/sativa/epac/version.py

#### Notes

- Several tools are listed without pinned versions in the Conda environments.
- When versions are not pinned, they are resolved at installation time and should be captured from the final runtime environment (for example, via conda list).

**Table S15:**
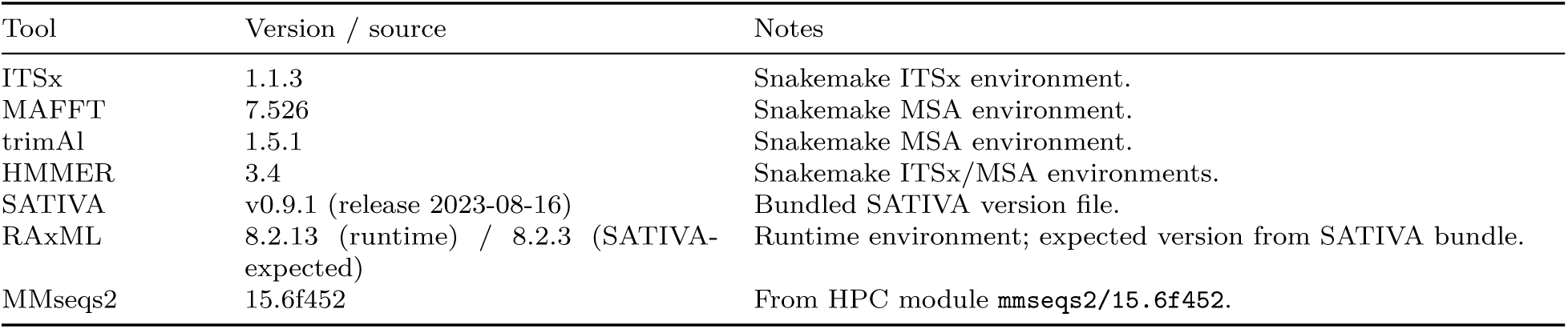
Configured software versions and provenance.

### C.2 B) ITSx parameters

Source: workflow/rules/itsx.smk and config/config.yaml.

Command template (per barcode):

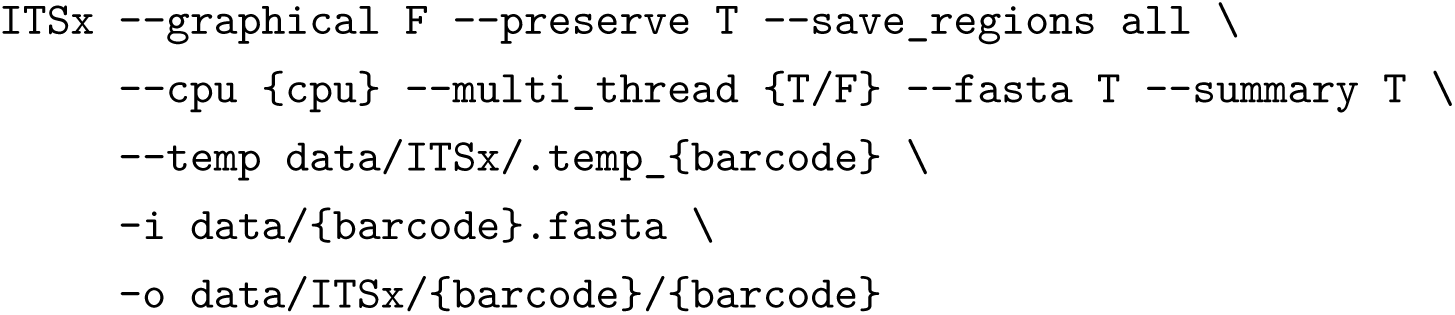

Deterministic mode:

- itsx_deterministic: true ⇒ –cpu 1, –multi_thread F
- Otherwise: –`cpu {itsx_cpu}` (default 1200 in `config.yaml`)

ITSx processing batch parameters (database side; from `config.yaml`):

- itsx.insert_batch_size: 10000
- itsx.status_update_chunk_size: 50
- itsx.parent_metadata_streaming: false
- itsx.parent_metadata_stream_fetch_size: 100
- itsx.parent_metadata_stream_filter_barcode: true
- itsx.parent_metadata_chunk_size: 50
- itsx.parent_metadata_fallback_chunk_size: 50
- itsx.parent_metadata_use_temp_table: false

### C.3 C) MAFFT parameters

Source: workflow/scripts/MSA/external_tools.py and config/config.yaml. Initial MSA:

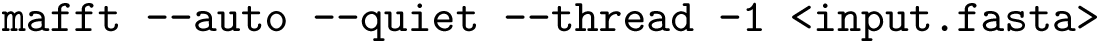

Adding sequences to an existing MSA:

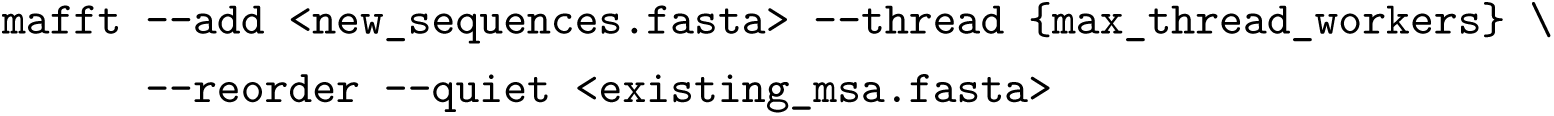

Threading configuration:

- msa.max_thread_workers: 50

### C.4 D) trimAl parameters

Source: workflow/scripts/MSA/external_tools.py and config/config.yaml.

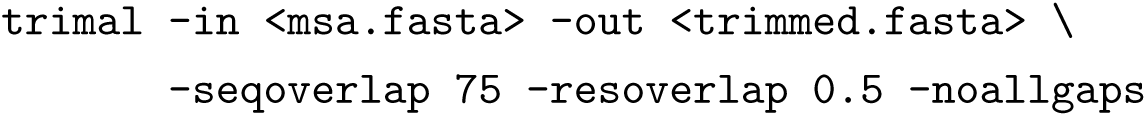

Thresholds:

- msa.trimal_seq_overlap: 75
- msa.trimal_res_overlap: 0.5

### C.5 E) HMMER parameters (hmmbuild/hmmscan/hmmpress)

Source: workflow/scripts/MSA/external_tools.py and config/config.yaml.

`hmmbuild:`

hmmbuild --dna -n {barcode}_{clade_name} <out.hmm> <msa.fasta>

`hmmpress:`

hmmpress -f <HMM_DB_PATH>

`hmmscan:`

`hmmscan --tblout <tbl.out> --cpu {threads} --noali <HMM_DB> <query.fasta>`

Assignment filtering:

- msa.classification_evalue_threshold: 1e-3 (applied after hmmscan in Python)
- msa.classification_max_workers: 20
- msa.classification_threads_per_worker: 4
- msa.classification_batch_size: 5000

### C.6 F) SATIVA parameters

Sources are the SATIVA workflow rule, the SATIVA wrapper script, and the central configuration file.

Command template (per MSA):

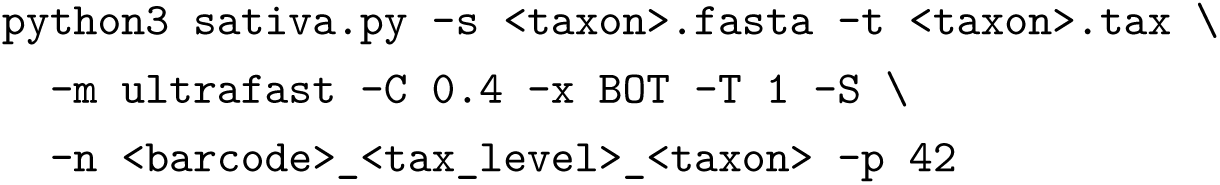

Key parameters:

- sativa.threads: 1
- sativa.sativa_mode: ultrafast
- sativa.confidence_threshold: 0.4
- sativa.taxonomy_code: BOT
- sativa.rand_seed: 42
- sativa.max_seq_count: 20000
- sativa.max_alignment_length: 5000
- sativa.timeout_seconds: 21600 (6 hours)
- sativa.tax_levels: family, genus, order, species

### C.7 G) MMseqs2 parameters

MMseqs2 clustering is run via an HPC job script (example at 99% identity):

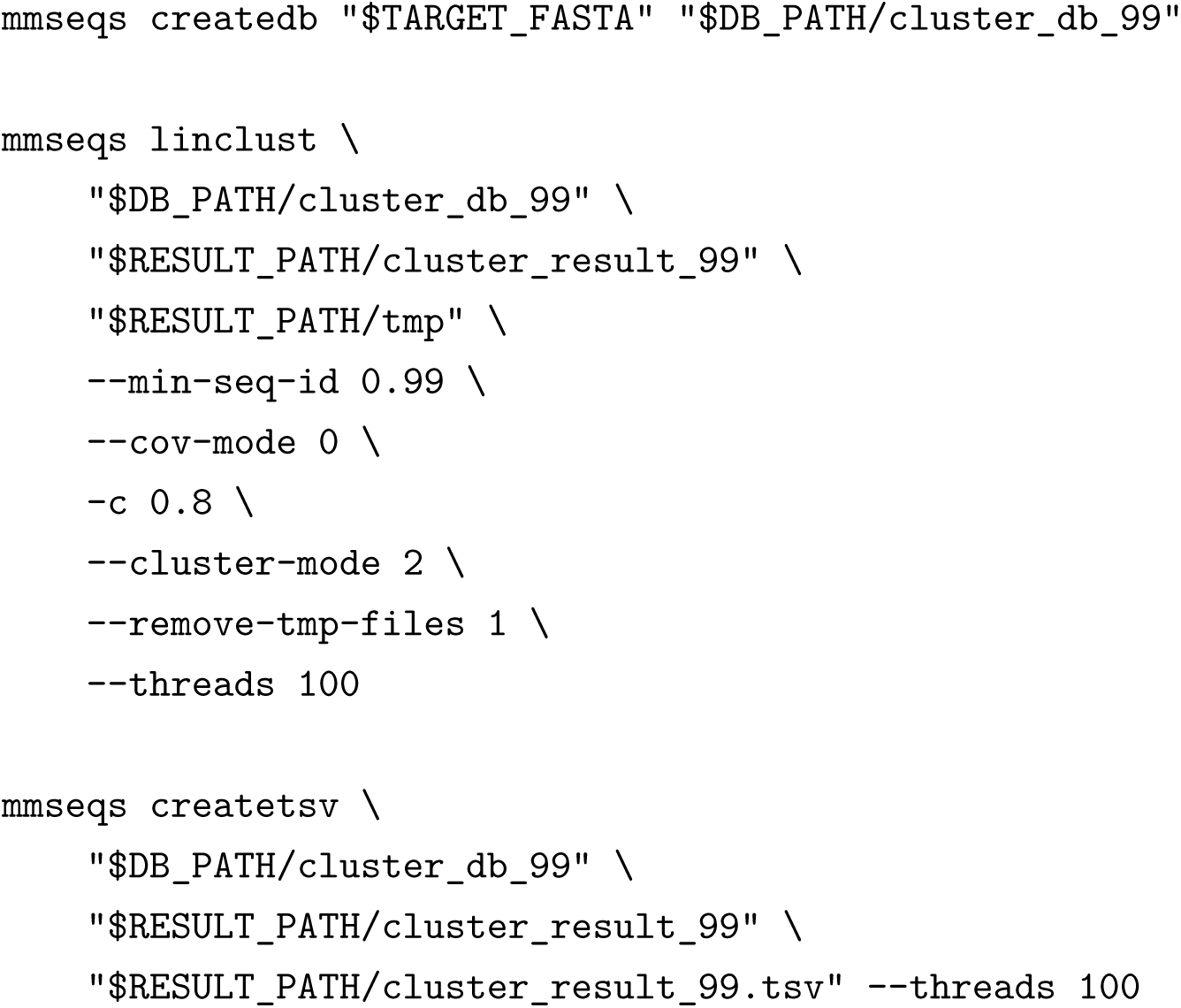

Notes:

- For 95% and 97% runs, –min-seq-id is set to 0.95 and 0.97 with the same parameters.
- Coverage mode 0 with -c 0.8 enforces 80% coverage on the shorter sequence.

**Table S16:**
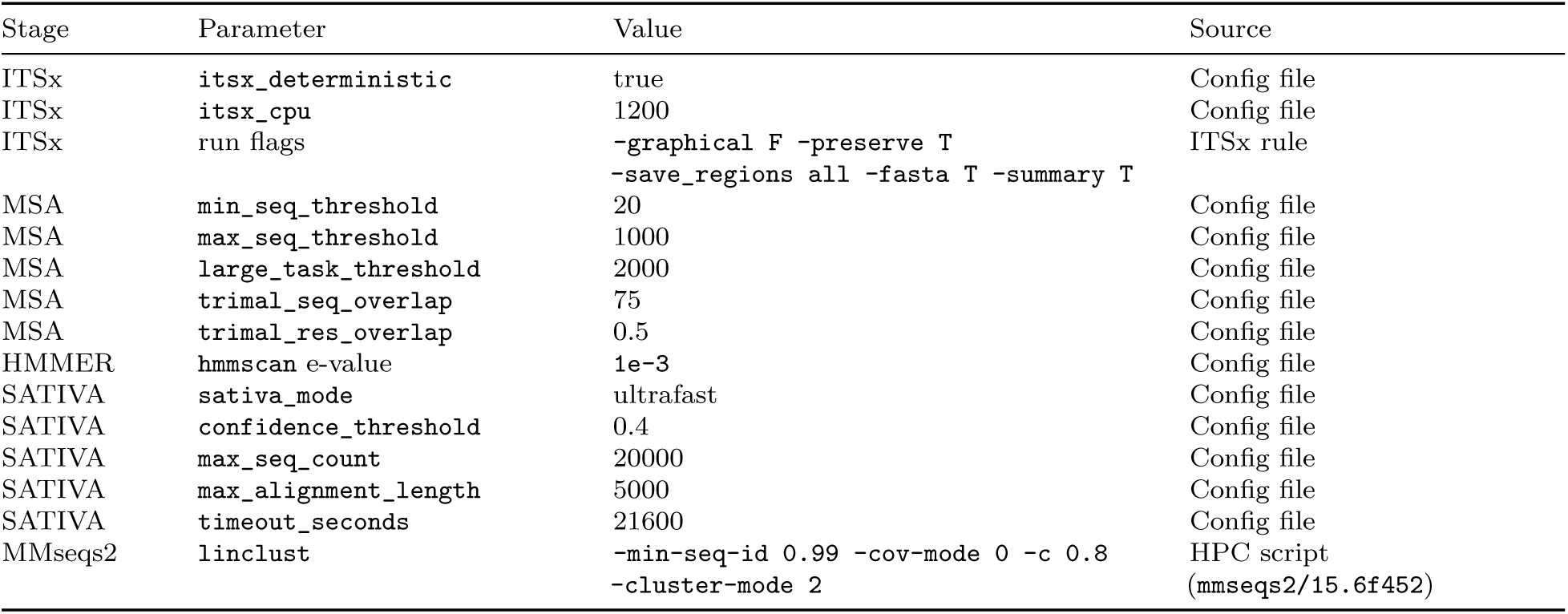
Key pipeline parameters and provenance.

### C.8 H) Table S1 — Key pipeline parameters (summary)

