## Supplementary for "CurateMake: an auditable workflow for multi-source ITS reference database harmonisation and phylogenetic validation"

#### A Reproducibility metadata

This appendix summarises the dataset versions, sequence counts, software versions, and key parameters used in the workflow. Dataset counts correspond to records retained after ingestion and pipeline-level filtering, rather than to raw download sizes. Software versions and parameters were extracted from the workflow configuration files, Conda environments, runtime modules, and pipeline scripts.

##### A.1 Data sources and dataset composition

All public reference datasets were extracted in late August 2025, although public release dates differed among sources (main-text Table ??). The full dataset was used for large-scale entropy analyses, whereas the public mini-dataset was used for demonstration, testing, and runtime benchmarking. Detailed barcode-level counts are provided in Table S1.

##### A.2 Controlled error-injection simulation (Experiment 2)

This section provides the reproducibility details of the controlled error-injection simulation summarised in the Methods and Results. The benchmark was designed to compare correction strategies under fully traceable ground truth rather than to characterise taxon-wide ecological performance.

**Table S1:** Barcode-level record counts after ingestion and pipeline filtering for the full and mini datasets. These counts are stratified by parsed barcode type and are not source-record totals: a source record can contribute more than one barcode-level record when multiple barcode annotations are retained.

| Dataset | Database | ITS | ITS1 | ITS2 | Total |
|---|---|---|---|---|---|
| Full | UNITE | 2,069,189 | 0 | 0 | 2,069,189 |
| Full | BOLD | 255,516 | 227,443 | 148,765 | 631,724 |
| Full | PLANiTS | 415,114 | 104,846 | 101,584 | 621,544 |
| Full | CALeDNA | 0 | 0 | 570,510 | 570,510 |
| Full | Total | 2,739,819 | 332,289 | 820,859 | 3,892,967 |
| Mini | UNITE | 1,034 | 0 | 0 | 1,034 |
| Mini | BOLD | 1,068 | 240 | 201 | 1,509 |
| Mini | PLANiTS | 1,959 | 406 | 405 | 2,770 |
| Mini | CALeDNA | 0 | 0 | 914 | 914 |
| Mini | Total | 4,061 | 646 | 1,520 | 6,227 |

##### A.2.1 Ground-truth anchor sequences

Twenty full-ITS sequences were manually retrieved from BOLD in January 2026 and used as ground-truth anchors: 10 plant species in *Carex* (Cyperaceae) and 10 fungal species in *Russula* (Russulaceae) (Table S2). These genera provide complementary test contexts for cross-kingdom contamination and within-genus misidentification. Species selection was restricted to entries with a CoL-accepted taxonomy and an unambiguous BOLD record, so that recovery could be evaluated against verifiable ground truth rather than confounded by nomenclatural ambiguity.

**Table S2:** Ground-truth anchor species used to build the controlled error-injection simulation. All species were confirmed as CoL-accepted at retrieval. **BOLD process ID** identifies the retrieved full-ITS record on BOLD; **GenBank** gives the corresponding INSDC nucleotide accession where available (a dash indicates records without a linked INSDC accession).

| *Carex* (Cyperaceae, Plantae) | | | *Russula* (Russulaceae, Fungi) | | |
|---|---|---|---|---|---|
| Species | BOLD process ID | GenBank | Species | BOLD process ID | GenBank |
| *C. annectens* | GBITS59088-21 | DQ115092 | *R. rhodocephala* | FUNCA3144-25 | PP870087 |
| *C. vulpinoidea* | GBITS43265-21 | FJ694692 | *R. fellea* | GBRUS538-13 | AM113958 |
| *C. aboriginum* | GBITS42856-21 | KT021126 | *R. azurea* | FABOL576-21 | – |
| *C. acaulis* | GBITS42857-21 | GU176147 | *R. dissimulans* | GBRUS1551-13 | HQ650756 |
| *C. acuta* | GBITS42858-21 | AF284992 | *R. granulata* | GBRUS1014-13 | EU598192 |
| *C. acutiformis* | GBITS59079-21 | HG915798 | *R. grisescens* | MQRUS1063-23 | PX308366 |
| *C. adrienii* | GBITS42859-21 | KP273628 | *R. gracillima* | NOBAS3287-16 | – |
| *C. alba* | GBITS42862-21 | AY278259 | *R. olivascens* | NOBAS484-15 | – |
| *C. angustata* | GBITS42865-21 | AF285015 | *R. pallescens* | NOBAS764-15 | – |
| *C. arcta* | GBITS59092-21 | DQ115098 | *R. pascua* | RADIS060-19 | MN992267 |

**Table S3:** Overall correction rate (%) by taxonomic corruption rate for the four correction strategies (mean ± SEM, $n = 5$ seeds), corresponding to **??**A. **CurateMake** (full pipeline) is the highest-performing strategy at every corruption level. SATIVA alone corrects nothing, and CoL-to-SATIVA is identical to CoL alone, confirming that phylogenetic validation contributes only within the workflow's alignment infrastructure.

| Corruption | Full pipeline | CoL alone | SATIVA alone | CoL-to-SATIVA |
|---|---|---|---|---|
| 1% | 43.4 ± 3.2 | 28.6 ± 3.9 | 0.0 ± 0.0 | 28.6 ± 3.9 |
| 2% | 38.6 ± 5.0 | 26.4 ± 3.6 | 0.0 ± 0.0 | 26.4 ± 3.6 |
| 3% | 44.6 ± 2.0 | 27.3 ± 1.9 | 0.0 ± 0.0 | 27.3 ± 1.9 |
| 5% | 39.4 ± 1.2 | 26.8 ± 1.7 | 0.0 ± 0.0 | 26.8 ± 1.7 |
| 10% | 42.5 ± 1.9 | 26.8 ± 0.8 | 0.0 ± 0.0 | 26.8 ± 0.8 |
| 15% | 42.4 ± 1.2 | 27.5 ± 1.0 | 0.0 ± 0.0 | 27.5 ± 1.0 |
| 25% | 41.7 ± 0.6 | 28.0 ± 0.6 | 0.0 ± 0.0 | 28.0 ± 0.6 |
| 50% | 42.6 ± 1.8 | 27.3 ± 0.2 | 0.0 ± 0.0 | 27.3 ± 0.2 |

**Table S4:** Per-error-type correction rate (%) at 15% corruption for the four strategies (mean ± SEM, $n = 5$ seeds), corresponding to **??**B, together with the preservation rate of non-corrupted records. `UNIDENTIFIABLE` errors are omitted (null correction by construction). CoL-to-SATIVA reproduces CoL alone exactly.

| Error type | Full pipeline | CoL alone | SATIVA alone | CoL-to-SATIVA |
|---|---|---|---|---|
| Obsolete taxonomy | 80.7 ± 2.9 | 67.5 ± 3.0 | 0.0 ± 0.0 | 67.5 ± 3.0 |
| Missing rank | 65.5 ± 1.8 | 42.7 ± 2.9 | 0.0 ± 0.0 | 42.7 ± 2.9 |
| Contamination (partial)[a] | 23.4 ± 6.4 | 0.0 ± 0.0 | 0.0 ± 0.0 | 0.0 ± 0.0 |
| at genus level | 23.1 ± 6.4 | 0.0 ± 0.0 | 0.0 ± 0.0 | 0.0 ± 0.0 |
| at kingdom level | 23.7 ± 6.4 | 0.0 ± 0.0 | 0.0 ± 0.0 | 0.0 ± 0.0 |
| Misidentification | 0.0 ± 0.0 | 0.0 ± 0.0 | 0.0 ± 0.0 | 0.0 ± 0.0 |
| Preservation (non-corrupted) | 99.8 ± 0.0 | 100.0 ± 0.0 | 100.0 ± 0.0 | 100.0 ± 0.0 |

[a] Full-rank contamination correction was negligible for all strategies; partial detection is the mean of the genus-level and kingdom-level recovery rates (reported on the two following lines).

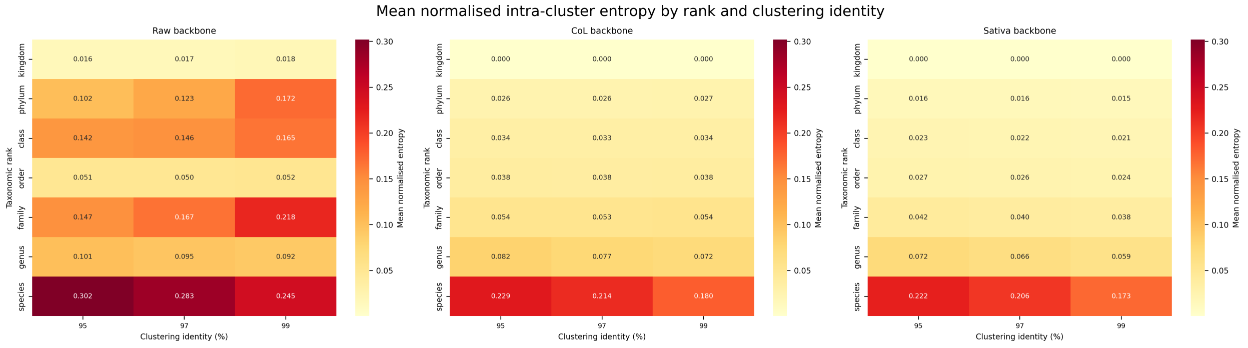


**Figure S1:** Mean normalised intra-cluster entropy ($H/\log_2 N$) per taxonomic rank and clustering identity threshold (95%, 97%, 99%), for each taxonomic annotation layer (Raw, CoL, SATIVA). Cell values are mean normalised entropy across all eligible non-singleton clusters. The species rank shows the highest residual entropy across all annotation layers and thresholds. The relative ordering of annotation layers (Raw > CoL > SATIVA) and the rank-entropy profile shape are qualitatively preserved across the three identity thresholds, supporting the robustness of the comparison presented in **??**.

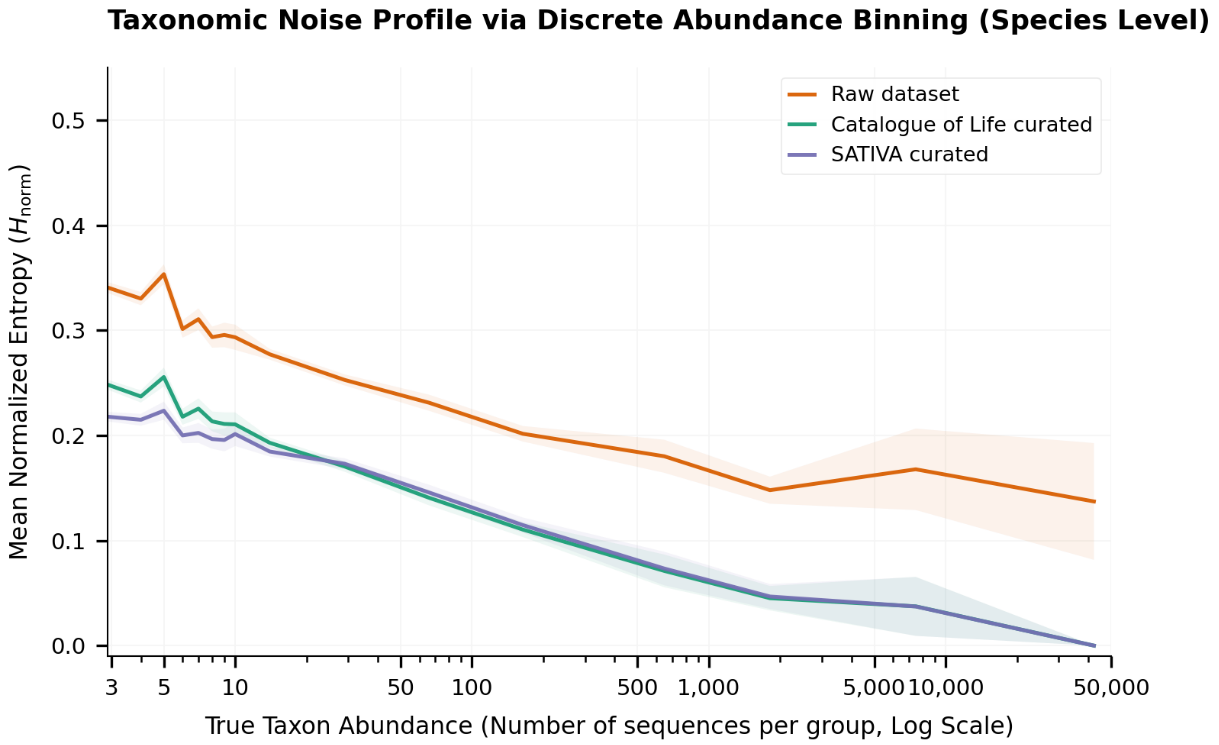


**Figure S2:** Mean normalised species-level intra-cluster entropy $(H/\log_2 N)$ as a function of true taxon abundance (number of sequences per taxon, log scale), for each taxonomic annotation layer (Raw, CoL, SATIVA). For all annotation layers, residual entropy decreases with taxon abundance: abundant taxa are curated to near-zero entropy, whereas rare taxa retain substantial heterogeneity. Most of the reduction relative to Raw is contributed by CoL harmonisation. SATIVA's additional reduction beyond CoL — the gap between the CoL and SATIVA curves — is concentrated at low abundance, where CoL leaves residual entropy; for abundant taxa, CoL already drives entropy close to zero, leaving little margin for further correction. This pattern is consistent with a floor effect (CoL saturating the achievable reduction in well-represented taxa) rather than a loss of phylogenetic signal in large groups. Entropy is measured at the species level, whereas MSA tasks are built at coarser ranks (family/genus, $\geq 20$ sequences): a rare species can therefore still receive a SATIVA species-level correction within a larger family- or genus-level alignment.

**Table S5:** Complete intra-cluster entropy comparison across annotation layers (Experiment 1). For each clustering identity threshold (95%, 97%, 99%) and taxonomic rank, columns **Raw**, **CoL**, and **SATIVA** give the mean normalised intra-cluster entropy ($H/\log_2 N$) across matched non-singleton clusters. The three rightmost columns give the Wilcoxon signed-rank test $p$-values for the pairwise layer comparisons (CoL vs Raw, CoL vs SATIVA, Raw vs SATIVA), computed on matched cluster entropies. $n$ is the number of matched non-singleton clusters at each threshold; this count is rank-independent because missing labels are retained as an "Unknown" category rather than excluded (see Methods). Values reported as $< 10^{-300}$ fell below double-precision floating-point resolution; "n.s." denotes non-significant comparisons ($p \geq 0.05$). Entropy decreases monotonically from Raw to CoL to SATIVA at every rank and threshold, and the only non-significant comparison is CoL vs SATIVA at kingdom level, consistent with CoL alone resolving most kingdom-level conflicts (see **??**).

| | Mean normalised entropy | | | Wilcoxon $p$-value | | |
|---|---|---|---|---|---|---|
| Rank | Raw | CoL | SATIVA | CoL vs Raw | CoL vs SATIVA | Raw vs SATIVA |
| *95% identity* ($n$ = 368,257 clusters) | | | | | | |
| Kingdom | 0.0163 | 0.0003 | 0.0003 | $< 10^{-300}$ | 0.077 (n.s.) | $< 10^{-300}$ |
| Phylum | 0.1019 | 0.0259 | 0.0163 | $< 10^{-300}$ | $< 10^{-300}$ | $< 10^{-300}$ |
| Class | 0.1418 | 0.0336 | 0.0231 | $< 10^{-300}$ | $< 10^{-300}$ | $< 10^{-300}$ |
| Order | 0.0507 | 0.0382 | 0.0273 | $< 10^{-300}$ | $< 10^{-300}$ | $< 10^{-300}$ |
| Family | 0.1469 | 0.0538 | 0.0418 | $< 10^{-300}$ | $< 10^{-300}$ | $< 10^{-300}$ |
| Genus | 0.1012 | 0.0819 | 0.0719 | $< 10^{-300}$ | $< 10^{-300}$ | $< 10^{-300}$ |
| Species | 0.3024 | 0.2290 | 0.2224 | $< 10^{-300}$ | $1.2 \times 10^{-243}$ | $< 10^{-300}$ |
| *97% identity* ($n$ = 400,109 clusters) | | | | | | |
| Kingdom | 0.0167 | 0.0003 | 0.0003 | $< 10^{-300}$ | 0.147 (n.s.) | $< 10^{-300}$ |
| Phylum | 0.1236 | 0.0260 | 0.0156 | $< 10^{-300}$ | $< 10^{-300}$ | $< 10^{-300}$ |
| Class | 0.1466 | 0.0333 | 0.0219 | $< 10^{-300}$ | $< 10^{-300}$ | $< 10^{-300}$ |
| Order | 0.0502 | 0.0377 | 0.0259 | $< 10^{-300}$ | $< 10^{-300}$ | $< 10^{-300}$ |
| Family | 0.1674 | 0.0535 | 0.0399 | $< 10^{-300}$ | $< 10^{-300}$ | $< 10^{-300}$ |
| Genus | 0.0956 | 0.0767 | 0.0657 | $< 10^{-300}$ | $< 10^{-300}$ | $< 10^{-300}$ |
| Species | 0.2839 | 0.2136 | 0.2068 | $< 10^{-300}$ | $1.0 \times 10^{-228}$ | $< 10^{-300}$ |
| *99% identity* ($n$ = 451,211 clusters) | | | | | | |
| Kingdom | 0.0180 | 0.0002 | 0.0002 | $< 10^{-300}$ | 0.211 (n.s.) | $< 10^{-300}$ |
| Phylum | 0.1726 | 0.0267 | 0.0150 | $< 10^{-300}$ | $< 10^{-300}$ | $< 10^{-300}$ |
| Class | 0.1656 | 0.0337 | 0.0205 | $< 10^{-300}$ | $< 10^{-300}$ | $< 10^{-300}$ |
| Order | 0.0519 | 0.0380 | 0.0243 | $< 10^{-300}$ | $< 10^{-300}$ | $< 10^{-300}$ |
| Family | 0.2182 | 0.0544 | 0.0379 | $< 10^{-300}$ | $< 10^{-300}$ | $< 10^{-300}$ |
| Genus | 0.0917 | 0.0721 | 0.0590 | $< 10^{-300}$ | $< 10^{-300}$ | $< 10^{-300}$ |
| Species | 0.2457 | 0.1797 | 0.1730 | $< 10^{-300}$ | $1.8 \times 10^{-168}$ | $< 10^{-300}$ |

**Table S6:** Taxonomic composition of the final harmonised database. For each rank, **Entries** is the number of barcode-resolved entries carrying a non-empty annotation at that rank, and **Taxa** is the number of distinct taxon names observed at that rank. Counts use the consolidated annotation per entry (SATIVA suggestion where available, otherwise CoL, otherwise Raw). The *Full database* columns cover all 5.19 M barcode-resolved entries (one entry per ITS/ITS1/ITS2 sub-region, so a full-length ITS sequence contributes several entries); the *Plants subset* columns are restricted to entries whose consolidated kingdom is Plantae. Entry counts decrease towards finer ranks as annotation becomes incomplete, while the number of distinct taxa increases. The species rank is the least complete: 2.4 M of 5.2 M entries (≈46%) in the full database and 613 K of 817 K (≈75%) in the plants subset are resolved to species.

| | Full database | | Plants subset | |
|---|---|---|---|---|
| Rank | Entries | Taxa | Entries | Taxa |
| Kingdom | 5.2 M | 7 | 817 K | 1 |
| Phylum | 5.0 M | 76 | 817 K | 9 |
| Class | 4.9 M | 201 | 796 K | 33 |
| Order | 4.4 M | 770 | 792 K | 185 |
| Family | 4.3 M | 2.6 K | 771 K | 694 |
| Genus | 4.1 M | 19.7 K | 745 K | 11 K |
| Species | 2.4 M | 183 K | 613 K | 92 K |

**Table S7:** Representative SATIVA correction examples from the controlled error simulation (seed 1, 15% corruption rate), under the full pipeline. Each row shows a sequence that was corrupted during the simulation, processed through CoL harmonisation, and then flagged and corrected by SATIVA. **Input taxonomy** is the label assigned after CoL harmonisation (before SATIVA); intermediate lineage ranks omitted with '. . . ' for clarity. **Correction level** is the highest rank at which SATIVA identified a mislabeling. **Proposed value** is SATIVA's suggestion for the mislabeled rank and all ranks below it. **Status**: ✓ = full agreement with ground-truth at that rank. All four examples are from the Carex/Russula simulation dataset; the underlying ITS sequences are Badread-generated synthetic reads from 20 ground-truth anchors. Seq. IDs are truncated to the first 8 characters.

| **Ex.** | **Error type** | **Input taxonomy (after CoL)** | **Correction level** | **Proposed value** | **Conf.** | **Status** |
|---|---|---|---|---|---|---|
| A | `CONTAMINATION` | *Plantae*; Tracheophyta; . . . ; Carex; Carex | Kingdom | Fungi | 0.999 | ✓[a] |
| B | `MISSING_RANK` | Fungi; Basidiomycota; Agaricomycetes; **Na; Na; Na** | Order | Russulales | 1.000 | ✓ |
| C | `MISSING_RANK` | Fungi; . . . ; Russulales; **Na; Na**; Russula | Family | Russulaceae | 0.999 | ✓ |
| D | `MISSING_RANK` | Fungi; . . . ; Russulaceae; **Na**; Russula azurea | Genus | Russula | 0.999 | ✓ |

[a] Contamination detection: the underlying ITS sequence originates from *Russula rhodocephala* (Fungi) but was relabeled as *Carex* (Plantae) by the corruption step; CoL validated the Plantae label; SATIVA correctly identifies the kingdom-level conflict from phylogenetic placement. Full proposed path: Fungi; Basidiomycota; Agaricomycetes; Russulales; Russulaceae; Russula; Russula (seq. 86481bf1, conf. 0.999).

**Table S8:** Distribution of SATIVA taxonomic suggestions on the full dataset, by suggestion type (`sativa_warnings.sativa_modification_type`). SATIVA produced 627,033 suggestions in total. **Completions** add one or more previously empty ranks below the validated lineage; **substitutions** replace an existing label at the deepest shared rank; **deletions** remove annotation and are split into single-rank (`DELETION_SIMPLE`) and multi-rank (`DELETION_MULTIPLE`) suggestions. Completions and substitutions are the most directly actionable classes; deletions are treated more cautiously because they remove existing annotation.

| Suggestion type | Count | % of total |
|---|---:|---:|
| Completion | 281,081 | 44.8 |
| Substitution | 171,552 | 27.4 |
| Deletion (total) | 174,400 | 27.8 |
| single-rank | 140,699 | 22.4 |
| multi-rank | 33,701 | 5.4 |
| Total | 627,033 | 100.0 |

**Table S9:** SATIVA suggestions on the full dataset by taxonomic rank of the correction (`sativa_warnings.mislabeled_level`) and suggestion type. Species is by far the most frequently corrected rank (334,273 suggestions, 53% of the total), consistent with SATIVA operating mostly on fine-rank labels. Completions dominate at intermediate ranks (phylum–family), whereas substitutions and deletions are concentrated at genus and species level.

| Rank | Completion | Substitution | Deletion | Total |
|---|---:|---:|---:|---:|
| Kingdom | 5 | 1,397 | 9 | 1,411 |
| Phylum | 27,150 | 2,599 | 10,703 | 40,452 |
| Class | 35,900 | 7,561 | 9,242 | 52,703 |
| Order | 7,203 | 13,155 | 3,355 | 23,713 |
| Family | 34,806 | 19,136 | 16,016 | 69,958 |
| Genus | 38,391 | 40,314 | 19,601 | 98,306 |
| Species | 137,626 | 84,843 | 111,804 | 334,273 |
| Other ranks[a] | 0 | 2,547 | 3,670 | 6,217 |
| Total | 281,081 | 171,552 | 174,400 | 627,033 |

[a] Intermediate ranks with few suggestions, pooled: domain, subclass, suborder, infraorder, superfamily, and tribe.

**Table S10:** SATIVA suggestions on the full dataset by barcode type and by source database, split by suggestion type. Counts are numbers of suggestions (`sativa_warnings`); deletions pool `DELETION_SIMPLE` and `DELETION_MULTIPLE`. Suggestions arise across all barcodes and all sources. UNITE, the largest source, contributes the most suggestions, and completions are the leading type for UNITE, BOLD, and PLANiTS, whereas the CALeDNA subsets — which enter with coarse "Eukaryota" kingdom labels — yield proportionally more substitutions.

| | Completion | Substitution | Deletion | Total |
|---|---|---|---|---|
| *By barcode* | | | | |
| ITS | 134,176 | 105,594 | 95,505 | 335,275 |
| ITS1 | 65,697 | 33,615 | 42,486 | 141,798 |
| ITS2 | 81,208 | 32,343 | 36,409 | 149,960 |
| *By source database* | | | | |
| UNITE | 157,303 | 70,802 | 99,925 | 328,030 |
| BOLD | 62,399 | 48,855 | 44,760 | 156,014 |
| PLANiTS | 52,292 | 11,385 | 9,326 | 73,003 |
| CALeDNA PITS | 2,624 | 14,975 | 6,760 | 24,359 |
| CALeDNA FITS | 6,463 | 25,535 | 13,629 | 45,627 |
| Total | 281,081 | 171,552 | 174,400 | 627,033 |

**Table S11:** Taxonomic modification rates per source, kingdom, and barcode for original sequences (`parent_seq_id IS NULL`, excluding duplicates). **pct_col_conflict**: proportion of sequences where CoL replaced at least one non-empty Raw rank with a different non-empty value (nomenclatural conflict). **pct_col_completiononly**: CoL changed the lineage but only by filling previously empty ranks (no conflict). **pct_sativa_conflict**: SATIVA replaced at least one non-empty CoL rank with a different non-empty value (phylogenetic re-assignment). **pct_sativa_mod**: any SATIVA modification (including rank completions by SATIVA). **pct_any**: final taxonomy (Sativa > CoL > Raw) differs from Raw at any rank. All rates are conservative lower bounds (pipeline-detected changes only); a conflict reflects a detected nomenclatural difference, not necessarily an annotation error. † CALeDNA systematically labels sequences as kingdom "Eukaryota" rather than the CoL-accepted "Fungi"/"Plantae"; the near-100% conflict rate reflects this labelling convention difference, not random misannotation.

| | n_seq | pct_col conflict | pct_col completiononly | pct_sativa conflict | pct_any |
|---|---|---|---|---|---|
| *By source* | | | | | |
| UNITE | 1,080,217 | 9.87 | 64.22 | 0.99 | 75.37 |
| BOLD | 387,703 | 13.41 | 85.29 | 9.51 | 98.70 |
| PLANiTS | 167,291 | 96.05 | 3.90 | 3.07 | 99.98 |
| CALeDNA PITS† | 122,153 | 99.45 | 0.55 | 8.75 | 100.0 |
| CALeDNA FITS† | 282,193 | 99.93 | 0.07 | 8.98 | 100.0 |
| *By kingdom (Raw label)* | | | | | |
| Fungi | 1,267,235 | 10.59 | 67.32 | 1.76 | 79.01 |
| Plantae | 340,406 | 52.69 | 47.29 | 8.71 | 99.99 |
| Animalia | 19,265 | 12.21 | 87.61 | 2.51 | 99.77 |
| Eukaryota† | 404,340 | 99.79 | 0.21 | 8.91 | 100.0 |
| Protista | 4,071 | 81.65 | 0.00 | 5.13 | 81.65 |
| *By barcode* | | | | | |
| ITS | 1,283,932 | 14.51 | 63.66 | 1.48 | 79.25 |
| ITS1 | 194,016 | 34.16 | 63.54 | 12.53 | 97.73 |
| ITS2 | 561,609 | 83.72 | 16.24 | 8.07 | 99.96 |

**Table S12:** ITSx extraction on the full dataset, by input barcode type (numerical values underlying the centre panel of **??**). **Input** is the number of deduplicated sequences of each barcode type entering ITSx; **ITS1** and **ITS2 extracted** are the sub-regions detected and re-inserted; **Reconstructed ITS** counts full ITS regions rebuilt from co-extracted ITS1/ITS2 sharing a parent; **No detection** is the number of sequences in which ITSx found no region (these are retained as pass-through records). Almost all extraction comes from full-length ITS inputs; pre-trimmed ITS1/ITS2 inputs rarely yield sub-regions because they lack the conserved flanking regions ITSx relies on.

| Input barcode | Input | ITS1 extracted | ITS2 extracted | Reconstructed ITS | No detection |
|---|---:|---:|---:|---:|---:|
| ITS | 1,692,277 | 1,674,716 | 1,667,949 | 1,655,388 | 4,037 |
| ITS1 | 231,697 | 191 | 44 | 39 | 231,209 |
| ITS2 | 607,844 | 68 | 147,832 | 56 | 386,176 |
| Total | 2,531,818 | 1,674,975 | 1,815,825 | 1,655,483 | 621,422 |

**Table S13:** Deduplication summary on the full dataset (100% identity over full-length sequences with identical taxonomy). The first pass, before ITSx, removes source-level redundancy; the second pass, after ITSx, removes duplicates introduced by sub-region extraction and reconstruction. **Duplicates removed by barcode** shows how the removed sequences of the second pass split across barcode types. The final 5,189,645 unique barcode-resolved entries constitute the harmonised database.

| Deduplication pass | Input | Unique kept | Duplicates removed |
|---|---:|---:|---:|
| First (pre-ITSx) | 3,581,943 | 2,536,799 | 1,045,144 |
| Second (post-ITSx) | 8,728,226 | 5,189,645 | 2,493,437 |
| *Second-pass duplicates removed, by barcode:* ITS 1,766,787 / ITS2 1,026,063 / ITS1 745,731 | | | |

**Table S14:** HMM-based barcode assignment on the full dataset (`hmmscan`, 2,918,862 sequences). Rows are the input barcode label at ingestion; columns are the barcode type assigned by the HMM library for alignment construction. Percentages are row-normalised (proportion of each input barcode type assigned to each output type). This step organises sequences with their close neighbours for alignment; it does not modify taxonomy. The high ITS1/ITS2 $\rightarrow$ ITS rates reflect inclusive matching of full-ITS profiles to sub-region queries (ITS1 and ITS2 are nested within the full ITS), not recovery of full-length signal; downstream analyses retain the original ingestion barcode.

| Input $\downarrow$ / Assigned $\rightarrow$ | ITS | ITS1 | ITS2 |
|---|---:|---:|---:|
| ITS | 1,352,134 (99.9%) | 584 (0.04%) | 558 (0.04%) |
| ITS1 | 339,755 (48.8%) | 310,075 (44.6%) | 45,739 (6.6%) |
| ITS2 | 555,666 (63.9%) | 43,443 (5.0%) | 270,908 (31.1%) |

### B Supplementary Figures

### C Software versions and parameters

This annex summarises tool versions (as configured) and the exact parameters used in the pipeline. Values are extracted from code and configuration files.

#### C.1 A) Software versions (as configured)

**Sources.**

- Conda environment files: `workflow/envs/*.yaml`
- SATIVA bundle: `workflow/scripts/Sativa/sativa/epac/version.py`

**Notes.**

- Several tools are listed without pinned versions in the Conda environments.
- When versions are not pinned, they are resolved at installation time and should be captured from the final runtime environment (for example, via `conda list`).

**Table S15:** Configured software versions and provenance.

| Tool | Version / source | Notes |
|---|---|---|
| ITSx | 1.1.3 | Snakemake ITSx environment. |
| MAFFT | 7.526 | Snakemake MSA environment. |
| trimAl | 1.5.1 | Snakemake MSA environment. |
| HMMER | 3.4 | Snakemake ITSx/MSA environments. |
| SATIVA | v0.9.1 (release 2023-08-16) | Bundled SATIVA version file. |
| RAxML | 8.2.13 (runtime) / 8.2.3 (SATIVA-expected) | Runtime environment; expected version from SATIVA bundle. |
| MMseqs2 | 15.6f452 | From HPC module `mmseqs2/15.6f452`. |

#### C.2 B) ITSx parameters

Source: `workflow/rules/itsx.smk` and `config/config.yaml`.

Command template (per barcode):

```
ITSx --graphical F --preserve T --save_regions all \
     --cpu {cpu} --multi_thread {T/F} --fasta T --summary T \
```

```
    --temp data/ITSx/.temp_{barcode} \
    -i data/{barcode}.fasta \
    -o data/ITSx/{barcode}/{barcode}
```

Deterministic mode:

- `itsx_deterministic: true` ⇒ `-cpu 1`, `-multi_thread F`
- Otherwise: `-cpu {itsx_cpu}` (default `1200` in `config.yaml`)

ITSx processing batch parameters (database side; from `config.yaml`):

- `itsx.insert_batch_size`: 10000
- `itsx.status_update_chunk_size`: 50
- `itsx.parent_metadata_streaming`: false
- `itsx.parent_metadata_stream_fetch_size`: 100
- `itsx.parent_metadata_stream_filter_barcode`: true
- `itsx.parent_metadata_chunk_size`: 50
- `itsx.parent_metadata_fallback_chunk_size`: 50
- `itsx.parent_metadata_use_temp_table`: false

#### C.3 C) MAFFT parameters

Source: `workflow/scripts/MSA/external_tools.py` and `config/config.yaml`.

Initial MSA:

```
mafft --auto --quiet --thread -1 <input.fasta>
```

Adding sequences to an existing MSA:

```
mafft --add <new_sequences.fasta> --thread {max_thread_workers} \
      --reorder --quiet <existing_msa.fasta>
```

Threading configuration:

- `msa.max_thread_workers`: 50

##### C.4 D) trimAl parameters

Source: `workflow/scripts/MSA/external_tools.py` and `config/config.yaml`.

```
trimal -in <msa.fasta> -out <trimmed.fasta> \
       -seqoverlap 75 -resoverlap 0.5 -noallgaps
```

Thresholds:

- `msa.trimal_seq_overlap`: 75
- `msa.trimal_res_overlap`: 0.5

##### C.5 E) HMMER parameters (hmmbuild/hmmscan/hmmpress)

Source: `workflow/scripts/MSA/external_tools.py` and `config/config.yaml`.

`hmmbuild`:

```
hmmbuild --dna -n {barcode}_{clade_name} <out.hmm> <msa.fasta>
```

`hmmpress`:

```
hmmpress -f <hmm_db_path>
```

`hmmscan`:

```
hmmscan --tblout <tbl.out> --cpu {threads} --noali <hmm_db> <query.fasta>
```

Assignment filtering:

- `msa.classification_evalue_threshold`: `1e-3` (applied after `hmmscan` in Python)
- `msa.classification_max_workers`: 20
- `msa.classification_threads_per_worker`: 4
- `msa.classification_batch_size`: 5000

##### C.6 F) SATIVA parameters

Sources are the SATIVA workflow rule, the SATIVA wrapper script, and the central configuration file.

Command template (per MSA):

```
python3 sativa.py -s <taxon>.fasta -t <taxon>.tax \
  -m ultrafast -C 0.4 -x BOT -T 1 -S \
  -n <barcode>_<tax_level>_<taxon> -p 42
```

Key parameters:

- `sativa.threads`: 1
- `sativa.sativa_mode`: ultrafast
- `sativa.confidence_threshold`: 0.4
- `sativa.taxonomy_code`: BOT
- `sativa.rand_seed`: 42
- `sativa.max_seq_count`: 20000
- `sativa.max_alignment_length`: 5000
- `sativa.timeout_seconds`: 21600 (6 hours)
- `sativa.tax_levels`: family, genus, order, species

##### C.7 G) MMseqs2 parameters

MMseqs2 clustering is run via an HPC job script (example at 99% identity):

```
mmseqs createdb "$TARGET_FASTA" "$DB_PATH/cluster_db_99"

mmseqs linclust \
    "$DB_PATH/cluster_db_99" \
    "$RESULT_PATH/cluster_result_99" \
    "$RESULT_PATH/tmp" \
    --min-seq-id 0.99 \
    --cov-mode 0 \
```

```
    -c 0.8 \
    --cluster-mode 2 \
    --remove-tmp-files 1 \
    --threads 100

mmseqs createtsv \
    "$DB_PATH/cluster_db_99" \
    "$RESULT_PATH/cluster_result_99" \
    "$RESULT_PATH/cluster_result_99.tsv" --threads 100
```

Notes:

- For 95% and 97% runs, `-min-seq-id` is set to 0.95 and 0.97 with the same parameters.
- Coverage mode 0 with `-c 0.8` enforces 80% coverage on the shorter sequence.

##### C.8 H) Table S1 — Key pipeline parameters (summary)

**Table S16:** Key pipeline parameters and provenance.

| Stage | Parameter | Value | Source |
|---|---|---|---|
| ITSx | `itsx_deterministic` | true | Config file |
| ITSx | `itsx_cpu` | 1200 | Config file |
| ITSx | run flags | `-graphical F -preserve T -save_regions all -fasta T -summary T` | ITSx rule |
| MSA | `min_seq_threshold` | 20 | Config file |
| MSA | `max_seq_threshold` | 1000 | Config file |
| MSA | `large_task_threshold` | 2000 | Config file |
| MSA | `trimal_seq_overlap` | 75 | Config file |
| MSA | `trimal_res_overlap` | 0.5 | Config file |
| HMMER | `hmmscan` e-value | `1e-3` | Config file |
| SATIVA | `sativa_mode` | ultrafast | Config file |
| SATIVA | `confidence_threshold` | 0.4 | Config file |
| SATIVA | `max_seq_count` | 20000 | Config file |
| SATIVA | `max_alignment_length` | 5000 | Config file |
| SATIVA | `timeout_seconds` | 21600 | Config file |
| MMseqs2 | `linclust` | `-min-seq-id 0.99 -cov-mode 0 -c 0.8 -cluster-mode 2` | HPC script (`mmseqs2/15.6f452`) |
